# SRC family kinase inhibition rescues molecular and behavioral phenotypes, but not protein interaction network dynamics, in a mouse model of Fragile X syndrome

**DOI:** 10.1101/2023.06.20.545800

**Authors:** Vera Stamenkovic, Jonathan D. Lautz, Felicia M. Harsh, Stephen E.P. Smith

## Abstract

Glutamatergic synapses encode information from extracellular inputs using dynamic protein interaction networks (PINs) that undergo widespread reorganization following synaptic activity, allowing cells to distinguish between signaling inputs and generate coordinated cellular responses. Here, we investigated how Fragile X Messenger Ribonucleoprotein (FMRP) deficiency disrupts signal transduction through a glutamatergic synapse PIN. In cultured cortical neurons or acute cortical slices from P7, P17 and P60 FMR1^-/y^ mice, the unstimulated protein interaction networks state resembled that of wildtype littermates stimulated with neurotransmitter agonists, demonstrating resting state pre-activation of signaling networks. We identified the Src family kinase (SFK) Fyn as a network hub, because many interactions involving Fyn were pre-activated. We tested whether targeting Fyn in FMR1^-/y^ mice could modify disease phenotypes, and found that Saracatinib (AZD-0530), an SFK inhibitor, normalized elevated basal protein synthesis, novel object recognition memory and social behavior in FMR1^-/y^ mice. However, SCB treatment did not normalize the PIN to a wild-type-like state *in vitro* or *in vivo*, but rather induced extensive changes to protein complexes containing Shank3, NMDARs and Fyn. We conclude that targeting abnormal nodes of a PIN can identify potential disease-modifying drugs, but behavioral rescue does not correlate with PIN normalization.

## Introduction

Protein-protein interactions among the hundreds of proteins mediating excitatory synaptic transmission are highly dynamic^1–3^. In neurons, protein complexes rapidly re-arrange their components in response to stimuli, which simultaneously changes the composition of the synapse and engages signal transduction machinery in a process basal to learning and memory^3–5^. Importantly, this glutamatergic synapse protein interaction network (gsPIN) is organized into discrete ‘modules’ of correlated protein interactions that change in unison in response to stimuli^4–7^. These modules may represent fundamental computational units through which neurons process information about the external environment and compute the appropriate cellular response to molecular inputs ^8, 9^.

Proteins encoded by genes linked to neurodevelopmental disorders (NDDs) comprise many nodes of the gsPIN^10–12^, which leads to the hypothesis that genetic mutations linked to NDDs may subtly alter how the gsPIN processes incoming signals^6, 13, 14^. Over developmental time, these alterations may lead to synaptic dysfunction, aberrant cellular or connectomic development of the brain, and the social and behavioral abnormalities associated with NDDs^15, 16^. We previously showed how two NDD-linked proteins that are physical network hubs of the gsPIN, Shank3 and Homer1, alter the modular gsPIN response to homeostatic plasticity^6^. In mice deficient in Shank3 or Homer1, the network state of the gsPIN in unmanipulated neurons resembled that of WT neurons undergoing homeostatic plasticity, leading to the hypothesis that the gsPIN was ‘pre-activated’ in knockout neurons. Indeed, following application of a stimulus that induced homeostatic plasticity in WT animals, pre-activated gsPIN networks lacking Shank3 or Homer1 failed to respond, because the state of many of their constituent interactions was already at a ceiling. This work^6^ supported the hypothesis that NDD-linked mutations may alter molecular computations performed by the gsPIN.

Here, we investigated the effect of another NDD-linked gene, FMR1 encoding Fragile X messenger ribonuclear protein (FMRP). FMRP is a polyribosome-associated RNA-binding protein involved in the transport and translational regulation of thousands of mRNA targets, many of which are involved in the development and maintenance of synapses^17, 18^. In humans, loss of FRMP expression causes Fragile X Syndrome (FXS), characterized by a complex phenotype consisting of intellectual disability, social anxiety, language deficits, hyperactivity, impulsivity, aggression, and disrupted sleep^19^. The lack of FMRP causes a molecular phenotype that is no less complex: excessive basal protein synthesis in neurons^20–22^, and a corresponding increase in protein turnover^23^, is associated with dendritic spines that are generally more abundant, less morphologically mature^24^, and hyperexcitable in response to glutamate uncaging^25^. A corresponding hyperexcitability of both metabotropic-^26, 27^ and NMDA-type receptor signaling^28^ leads to deficits in synaptic plasticity, most notably mGluR-dependent long term depression (LTD)^27^, and may contribute to cortex-wide hyperactivity on electroencephlogram in both human^29, 30^ and FMR1 knockout (FMR1^-/y^) mice^31^. Intriguingly, mGluR-dependent LTD is mTOR-dependent in WT but not FMR1^-/y^ neurons, suggesting a fundamental re-wiring of signal transduction^32^. Since FMRP is involved in the activity-dependent regulation of mRNA translation of many physical members of the gsPIN, we predicted that we could directly measure changes in not only the composition of the gsPIN, but also the way in which the gsPIN processes incoming information in FMR1^-/y^ neurons.

We stimulated cultured cortical neurons or acute cortical slices with agonists of NMDA or metabotropic glutamate receptors and characterized genotype- and activity dependent changes in gsPIN interactions. Protein network analysis demonstrated basal network hyperactivity in FMR1^-/y^ mice that was consistent with both mGluR5 and NMDAR-mediated hyperactivity, and that included changes in protein complexes containing the kinase Fyn. Inhibition of Fyn with saracatinib (SCB), an FDA-phase-2 kinase inhibitor that targets Fyn and other SRC-family kinases (SFK), rescued elevated basal protein synthesis, learning and memory deficits, and social behavior in FMR1^-/y^ mice. However, SFK inhibition did not normalize the gsPIN to a ‘wild-type-like’ state, but rather modified it by inducing changes in protein interactions containing Shank3, NMDARs and Fyn. Thus, identifying and targeting abnormal nodes of the gsPIN can suggest potential targets for disease-modifying drugs, but even effective drug modification may not normalize the underlying pathology.

## Results

### Altered protein interaction network dynamics following neurotransmitter agonist exposure in FMR1^-/y^ cultured neurons

To test if FMR1 mutations disrupt signal transduction through the gsPIN, we monitored activity-dependent synaptic protein interactions after stimulation of cultured cortical neurons from FMR1^-/y^ mice or WT controls with NMDA/glycine (to activate NMDA receptors), DHPG (to activate type 1 metabotropic glutamate receptors, mGluR1/5), or artificial cerebrospinal fluid (aCSF, a negative control)^5^. Importantly, we included antagonists for non-targeted receptors, including the NMDAR antagonist APV in the DHPG stimulation buffer^28^. Following 5 minutes of acute treatment, the timepoint of maximum module activation^7^, we performed quantitative multiplex co-immunoprecipitation (QMI), an ELISA-like antibody-based flow cytometry assay that performs hundreds of simultaneous co-immunoprecipitations on cell lysates to measure protein interactions that acutely change following a signal transduction event^33^ (Fig 1A). Following transformation of QMI data matrices into principal component space, WT cultured neurons showed a typical^5, 7^ pattern of activation: NMDA produced a strong response in principal component 1, while DHPG produced an orthogonal response in principal component 2 (Fig 1B). Similarly, correlation network analysis (CNA), which clusters interactions based on similar patterns of changes across experiments^34^, clustered protein interactions into 3 discrete modules: one that correlated with NMDA treatment and one that correlated DHPG treatment, as well as one that correlated with genotype (Fig 1, C,D). These data recapitulate previously published work showing that NMDA and DHPG activate independent modules of correlated protein-protein interactions in WT neurons^5, 7^.

**Figure 1:**
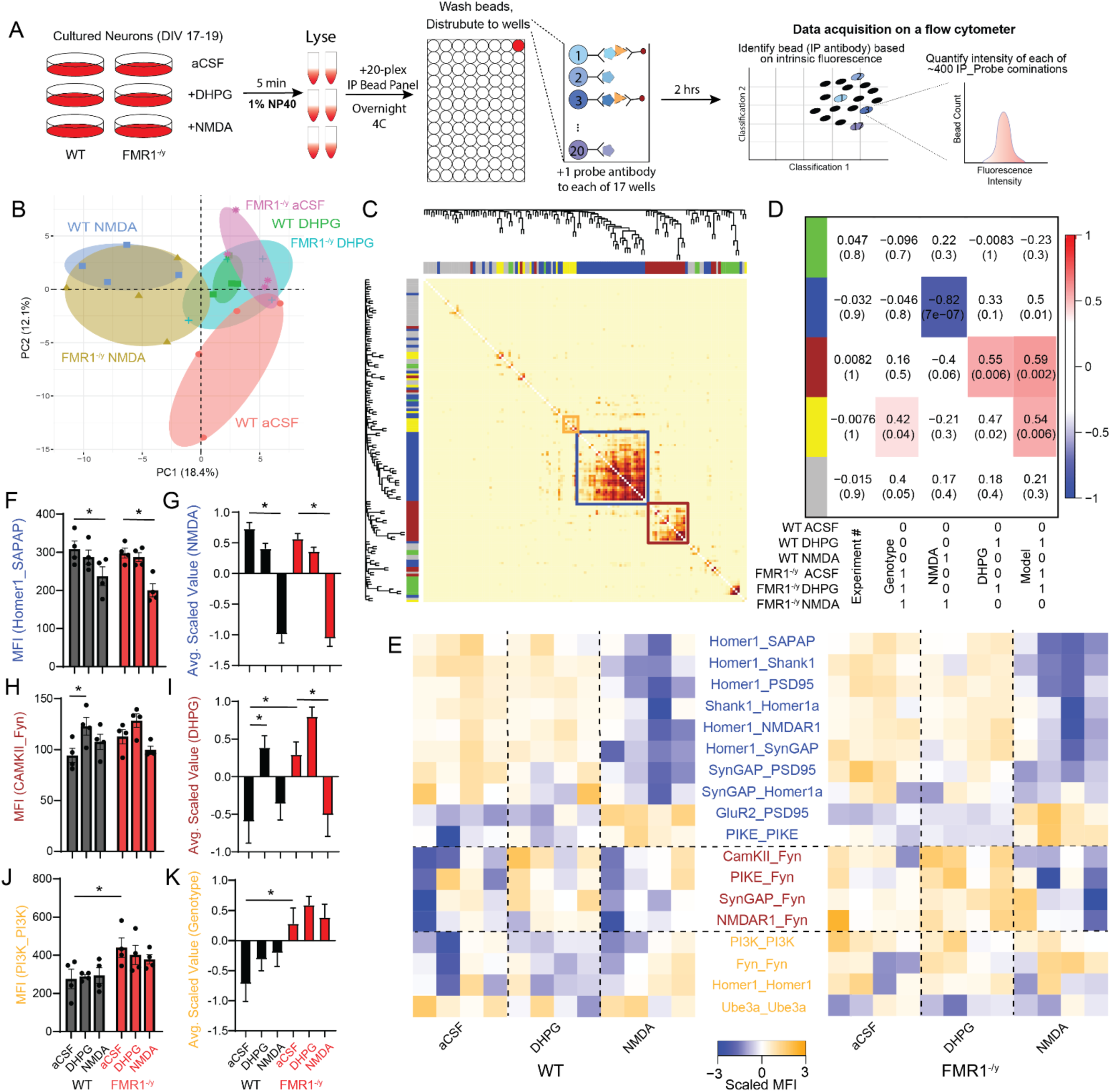
FMR1^-/y^ cultured cortical neurons show tonic activation of DHPG-responsive protein interactions. A) Experimental design. B) Principal component graph of median fluorescent intensity (MFI) matrices from N=4 cultures of FMR1^-/y^ or WT neurons treated with aCSF, NMDA or DHPG. Each colored dot represents an independent agonist-treated cell culture. C) Topological overlap matrix (TOM) plot showing clustering of individual protein interactions into modules. Each point on the dendrograms represents an individual interaction, each pixel of the plot indicates the correlation between the interaction on the X and Y axis (dark = higher correlation coefficient), and the colored boxes outline each module of interest. D) Module-trait correlation table shows the correlation coefficient (top number) and p-value (bottom number) of the correlation between the eigenvector describing each module (indicated by colored bars at left), and the binary-coded hypothesis listed below the table. E) Interactions that were both individually statistically significant by a Bonferroni-corrected adaptive non-parametric test corrected for multiple comparisons comparing treatment groups (ANC, see methods), and were a member of a module of interest, are represented by a row-scaled heatmap (blue-low abundance, orange-high abundance). Interaction labels are colored based on their assigned module. F) Example of an interaction in the NMDA-responsive blue module, Homer1_SAPAP. MFI, median fluorescent intensity. Asterisk represents statistically significant difference measured by ANC. G) Averaged scaled value of all interactions in the NMDA-responsive blue module. Asterisks represent statistically significant difference by one-way ANOVA followed by Sidak’s multiple comparison test, p<0.05. H) Example of an interaction in the DHPG-responsive brown module, CAMKII_Fyn. Asterisk represents statistically significant difference by ANC, p<0.05. I) Averaged scaled value of all interactions in the DHPG-responsive brown module. Asterisks represent statistically significant difference by one-way ANOVA followed by Sidak’s multiple comparison test, p<0.05. J) Example of an interaction in the genotype-correlated yellow module, PI3K_PI3K. Asterisk represents statistically significant difference by ANC, p<0.05. K) Averaged scaled value of all interactions in the genotype-correlated yellow module. Asterisks represent statistically significant difference by one-way ANOVA followed by Sidak’s multiple comparison test, p<0.05. N = 4 cultures from 4 different mice per condition.

In FMR1^-/y^ neurons, NMDA-treated samples clustered with WT-NMDA samples in PCA space. However, FMR1^-/y^ aCSF-treated neurons clustered with WT neurons treated with DHPG, and DHPG stimulation produced no additional changes (Fig 1B). This pattern suggests that FMR1^-/y^ neurons show basal hyperactivation of DHPG-like signaling on the level of the gsPIN. Moreover, the eigenvector describing the DHPG-responsive module identified by CNA correlated with a vector model in which FMR1^-/y^-aCSF was equivalent to both FMR1^-/y^-DHPG and WT-DHPG (Fig 1D, “Model”). These data support a hypothesis that protein interaction network states that change in response to mGluR agonism in WT mice are already activated at baseline in FMR1^-/y^ mice, consistent with the mGluR5 hyperactivity hypothesis of Fragile X^21, 26, 35^.

We identified high-confidence protein-protein interactions that changed in response to NMDA or DHPG by identifying interactions that both (1) met membership criteria for one of the three CNA modules of interest, and (2) that were statistically significant in paired, binary comparisons of aCSF vs. DHPG or aCSF vs. NMDA within-genotype, or WT-aCSF vs. FMR1^-/y^ -aCSF, using a multiple-comparison-corrected statistical test designed for QMI data^36^. The NMDA-responsive blue module followed the established pattern^5, 7^ of a strong reduction in the abundance of interactions containing synaptic scaffold proteins Homer1, Shank1 and PSD-95, as well as signaling proteins SynGAP and SAPAP (Fig 1E). The interaction IP Homer, probe SAPAP (represented as Homer1_SAPAP) most strongly correlated with the eigenvector describing the module, and is shown as a typical example (Fig 1F). Additionally, GluR2_PSD95 and PIKE-PIKE increased with NMDA treatment, indicative of increased synaptic recruitment of AMPA receptors and PI3K pathway activation. The behavior of a module is often more consistent across samples than the individual interactions that comprise the module, since biological noise inherent in signal transduction systems averages out over a larger number of measurements^7^, so we calculated the average scaled absolute value of all interactions in the NMDA-responsive module. We observed a significant reduction in average scaled intensity in response to NMDA in both genotypes (Fig 1G). For the DHPG-responsive brown module, only 4 interactions met the stringent double-significant criteria, all which involved the SRC family kinase Fyn (e.g. CAMKII_Fyn, Fig 1H), which suggests that Fyn activation plays a key role in DHPG-dependent signaling. In FMR1^-/y^ neurons, the average scaled absolute value of the DHPG-responsive module was already elevated in aCSF, and DHPG treatment failed to elicit additional module activation (Fig 1I). A yellow, genotype-dependent module contained interactions such as PI3K_PI3K (Fig 1J) and Fyn_Fyn, and was increased in FMR1^-/y^ neurons regardless of treatment (Fig 1K). Overall, these data support the hypothesis that in FMR1^-/y^ neurons, protein interaction networks downstream of NMDA stimulation are intact, but those downstream of mGluR5 stimulation are in a ‘pre-activated’ state, characterized by tonically elevated protein interactions involving the kinase Fyn.

### Altered protein interaction network dynamics following neurotransmitter agonist exposure in FMR1^-/y^ acute brain slices

We next investigated protein network alterations downstream of mGluR5 and NMDA receptor agonism in acute brain slices from WT and FMR1^-/y^ mice. Since responses may change in both WT and FMR1^-/y^ animals as neurons mature, we investigated three different ages of mice, postnatal day (P) 7 (approximately equivalent to a human infant), P17 (adolescence), and P60 (adult), and analyzed each age independently, as above (Fig 2A and Fig S1-S3). At each age, we identified a module that responded preferentially to NMDA, and one that responded more strongly to DHPG in WT animals. Although the modules were calculated independently at each age, and often consisted of different interactions, we color-coded them as NMDA-blue and DHPG-brown to highlight the consistent biological response. In the WT condition at all ages, the NMDA module had a high average scaled value in aCSF, and showed a strong negative change in NMDA (Fig 2B-D). At P17, the NMDA module also responded to DHPG with a significant reduction in scaled intensity. In FMR1^-/y^ slices in aCSF, the NMDA module also had a high average scaled value, not significantly different from WT_aCSF, and also showed a strong negative response to NMDA. Unlike WT animals, the NMDA module responded to DHPG at both P17 and P60, and showed a trend towards a response to DHPG at P7. These NMDA module summary data are consistent with node-edge diagrams of all interactions that changed significantly in response to NMDA in both genotypes across all ages (Fig S4). In WT animals, the magnitude of the NMDA response peaked at P17 before becoming much less intense at P60, likely reflecting network stabilization with maturation. FMR1-/y slices showed a similar developmental trajectory to WT, and a similar magnitude of response to NMDA (although note that many mGluR5-containing interactions were dysregulated in the P60 FMR1-/y slices) (Fig S4). Overall, NMDA module dynamics appeared intact in FMR1^-/y^ mice, but the module responded unusually strongly DHPG.

**Figure 2.**
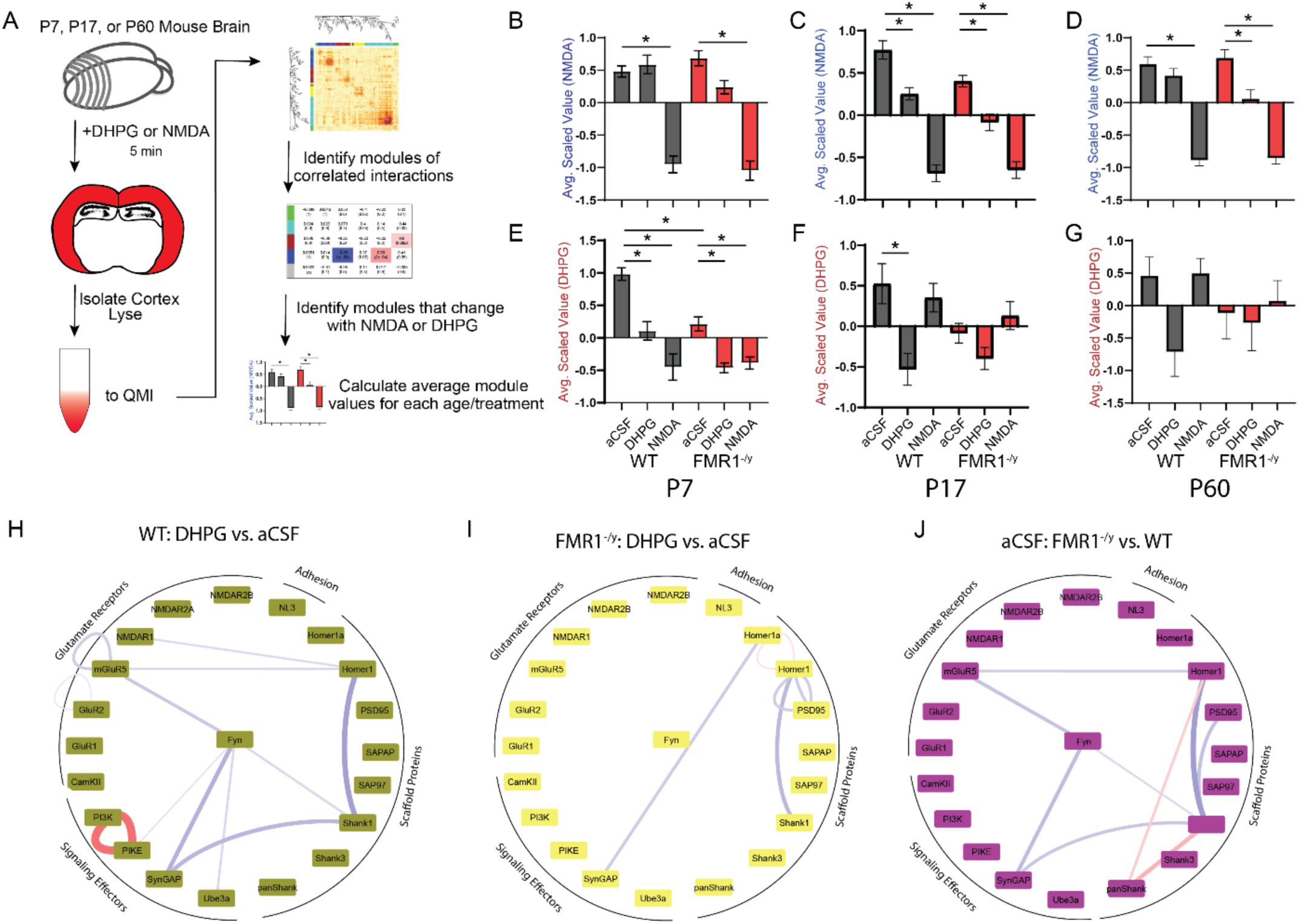
Hyperactive tonic DHPG-like signaling throughout development in FMR1^-/y^ brain slices. **A)** Experimental design. B-G) Average scaled values of protein interaction modules that respond to NMDA (blue, B-D) and DHPG (brown, E-G) in WT and FMR1**^-/y^** brain slices at P7 (B,E), P17 (C,F) and P60 (D,G). Asterisks represent statistically significant difference by one-way ANOVA followed by Sidak’s multiple comparison test, p<0.05. H-J) Node-edge diagrams show protein interactions that changed in response to DHPG stimulation (compared to aCSF) in WT (H) or FMR1^-/y^ slices (I) at P17, or interactions that differed comparing FMR1^-/y^_aCSF vs. WT_aCSF at p17 (J). Nodes represent proteins, edges represent protein-protein interactions that were significantly different in the given comparison; red = increased, blue = decreased. The relative magnitude of the change is reflected in edge thickness. All interactions shown were both statistically significant by a Bonferroni-corrected adaptive non-parametric test corrected for multiple comparisons (ANC), and were a member of a significant module. N = 4 animals per age/condition.

The DHPG-responsive “brown” module also showed a high relative scaled value in WT_aCSF, and consistently showed a strong negative change in WT_DHPG at all ages (Fig 2E-G). At P7, the module also responded to NMDA (Fig 2E), but no longer responded to NMDA at later ages. In FMR1^-/y^ untreated (aCSF) slices, the module was consistently reduced compared to WT_aCSF, suggesting basal network hyperactivity (Fig 2E-G). This basal reduction in module interactions resulted in the module no longer distinguishing aCSF, DHPG and NMDA stimulation. For example, in P7 WT slices, the module responded to both DHPG with a moderate reduction, and NMDA with a stronger reduction. In contrast, in FMR1^-/y^ slices, DHPG and NMDA both resulted in an equally strong dissociation. In P17 and P60 animals, the basal reduction of the DHPG module in FMR1^-/y^ slices led to a lack of change in response to DHPG. This ‘pre-activation’ was also evident in node-edge diagrams of all interactions that changed significantly in response to DHPG (Fig S5). In WT animals, the magnitude of DHPG signaling peaked at P17, while P60 slices showed a very small response. In FMR1^-/y^ animals, the magnitude of the difference between DHPG and aCSF was smaller (FigS5, middle column), but the difference between FMR1_aCSF and WT_aCSF was similar to DHPG treatment (Fig S5, right), supportive of the hypothesis that mGluR5 signaling is ‘pre-activated’. The developmental evolution of the DHPG response was similar in both genotypes. Overall, basal pre-activation of mGluR signaling in FMR1^-/y^ slices was consistent across ages, despite differences in the protein interactions that comprise the modules.

### SCB treatment rescues molecular and behavioral phenotypes in FMR1**^-/y^** mice

We next aimed to manipulate this ‘pre-activated’ protein network during development, to ask if reducing tonic signaling could modify disease phenotypes. Since direct antagonism of mGluR5 signaling has not performed well clinically^37^, we hypothesized that Fyn kinase may be a promising target. Fyn physically associates mGluR5^38^, and mGluR5 activation by DHPG or Amyloid Beta leads to Fyn activation^38^. Activated Fyn phosphorylates mGluR5, increasing its synaptic recruitment^39^, and phosphorylates NMDA receptors, reducing their co-association with PSD95^40^. We noticed that Fyn-containing protein interactions mediated the DHPG response in cultured cortical neurons (Fig 1), and that Fyn interactions were already in an “activated” state in aCSF conditions in P17 FMR1^-/y^ slices, and not changed after DHPG stimulation (Fig 2, H-J). Thus, we hypothesized that inhibition of hyperactive Fyn kinase during mid-development may de-activate the tonically active protein network. We identified a small molecule, Saracatinib (SCB), that inhibits brain-expressed SFKs including Fyn, Src, Lyn and Yes with an IC_50_ in the low nanomolar range (due to high levels of homology among SFKs, specific inhibitors of Fyn kinase are not available). SCB has been successfully used to treat preclinical animal models of Parkinson’s disease, Alzheimers disease and epilepsy^41–48^. SCB is orally bioavailable, crosses the blood-brain-barrier, and was safe and tolerated in a human phase 2 clinical trial for Alzheimer’s disease^49^, making it an attractive translational candidate.

Excessive protein synthesis is a robust phenotype present in both FMRP-deficient mouse brain and human iPS neurons^20^ that is easily measured by the SUNSET assay^50^. To test if SCB normalizes excess protein synthesis in FMR1^-/y^ mice, we incubated acute cortical slices with puromycin, in the presence or absence of SCB (Fig 3A). SCB reduced SFK phosphorylation in both genotypes (Fig 3B,C). In FMR1^-/y^ untreated slices, puromycin incorporation was increased, reflecting elevated protein synthesis. SCB normalized protein synthesis rates to WT levels, but did not affect WT protein synthesis rates (Fig. 3B,D).

**Figure 3.**
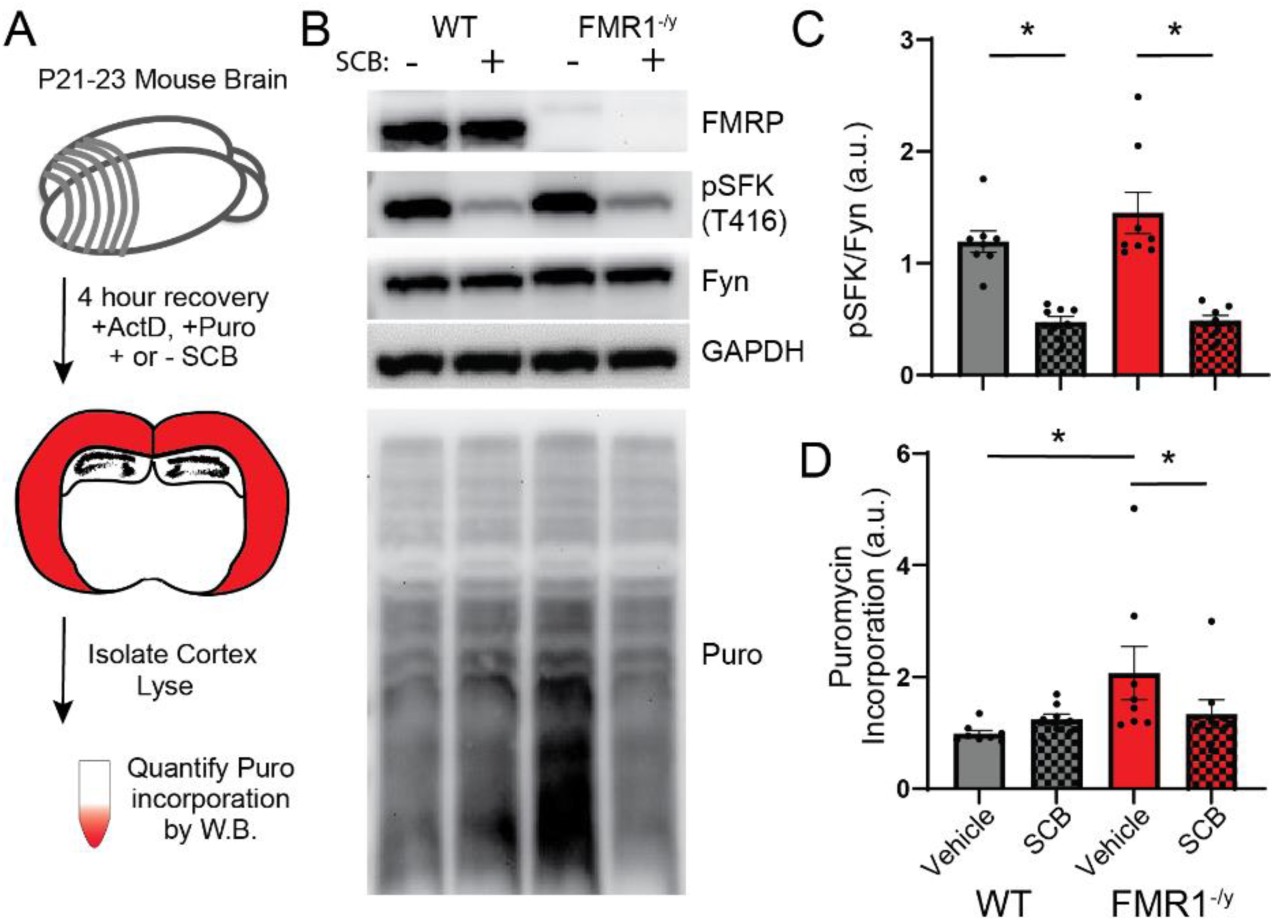
Pharmacological reduction of SFK activity restores protein synthesis rates in FMR1^-/y^ acute cortical slices. **A)** Experimental design. B) Representative Western blots. C) SCB reduced SFK phosphorylation in WT and FMR1^-/y^ slices. D) SCB significantly lowered puromycin incorporation in FMR1^-/y^ slices without altering its levels in WT. * p < 0.05 by one-way ANOVA followed by Sidak’s multiple comparison test. N= 8 animals per condition (3 slices per animal).

We next tested the ability of SCB to rescue FXS-related mouse behavior by treating FMR1^-/y^ mice, or their WT littermates, with SCB beginning after weaning (P23-26). Importantly, prior studies identified dosing schedules that result in drug levels in the mouse brain that are comparable to those achieved in human CSF during phase 2 clinical trials^42^, so we used an established oral gavage dosing strategy that achieves steady-state clinically relevant levels of SFK inhibition in the brain. Administration of the drug or vehicle began seven days prior to testing and continued for the duration of the behavioral testing (Fig. 4A). To test social behavior, we used a three-chamber social interaction test in which WT mice show an innate preference for spending time in a chamber containing a caged mouse, compared to an empty container^51^. Unlike WT controls, vehicle-treated FMR1^-/y^ mice did not show a significant preference for spending time in the social chamber (Fig 4B) or for interacting with a caged conspecific (Fig 4C), but SCB-treated FMR1^-/y^ mice showed a significant preference for both social measures. The mice showed no innate preferences for either side of the apparatus during acclimation (Fig S6A). To test learning and memory, we used a novel object recognition test in which WT mice prefer to explore a novel object compared to an object they were exposed to 48 hours prior. Unlike WT controls, vehicle-treated FMR1^-/y^ mice did not show a preference for exploring the novel object, but SCB-treated FMR1^-/y^ mice spent significantly more time investigating the novel compared to the familiar object (Fig. 4D). The mice did not show an innate preference for either object in the training phase (Fig S6B). In the open field test, which measures locomotion and anxiety-like behavior, mice showed no differences between groups for total distance traveled, time spent in the center, or entries into the center (Fig S6C-F). Repetitive grooming, a stereotyped mouse behavior that may reflect the increased rate of stereotypies in children with Fragile X and autism, was elevated in FMR1^-/y^ mice regardless of SCB treatment (Fig 4E). Thus, SCB treatment in juvenile FMR1^-/y^ mice improved social interactions and memory, but did not improve repetitive grooming.

**Fig. 4.**
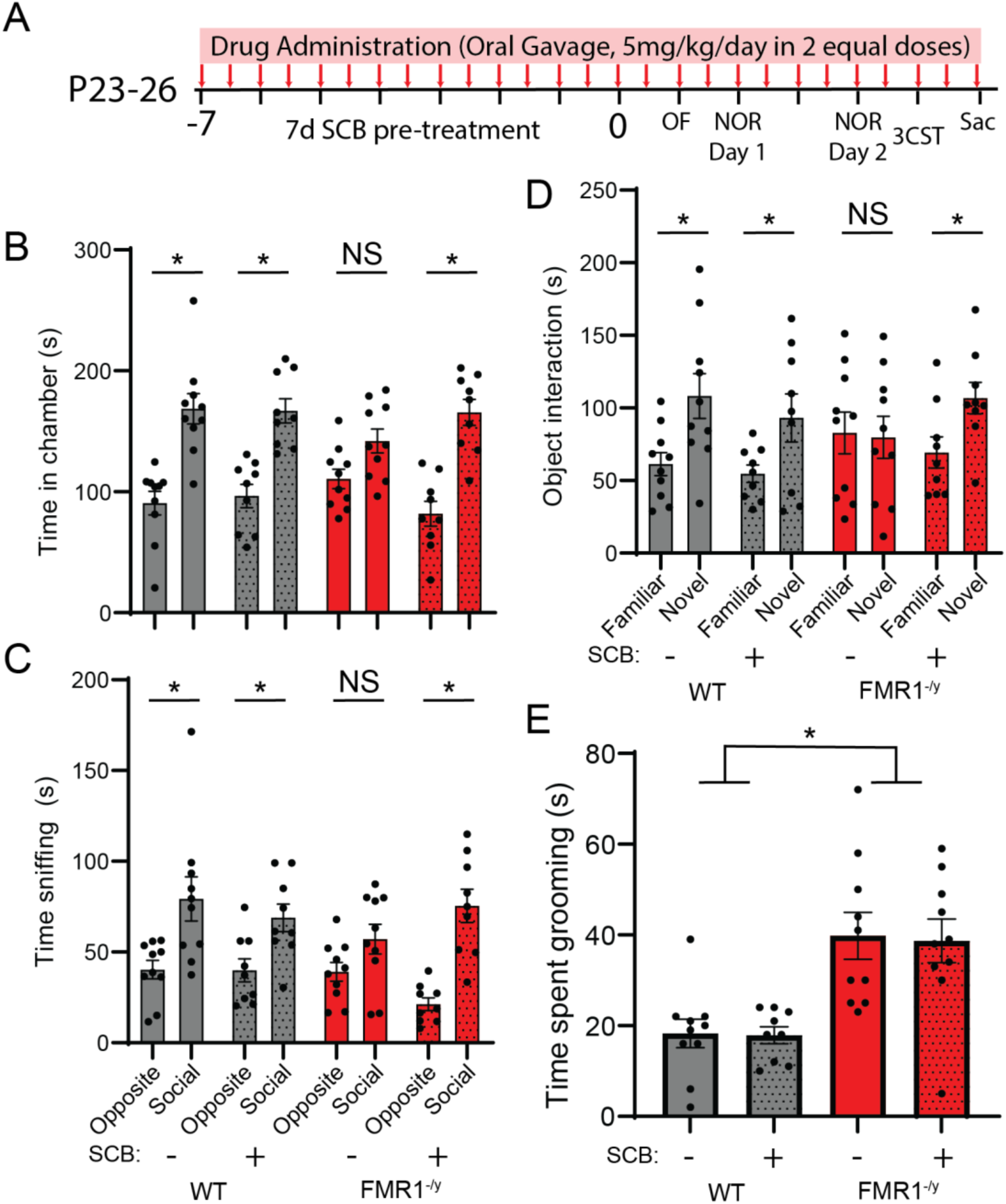
SCB treatment rescues impaired social behavior and object recognition memory in FMR1^-/y^ mice. **A)** Experimental design. B) Time spent in the social vs. opposite chamber and C) time spent sniffing the social-mouse enclosure vs. the empty enclosure in the three-chamber social test (3CST). p<0.05 by t-test comparing the social vs. opposite chamber of each treatment/genotype. D) Time spent interacting with the novel vs. familiar object on day 2 of the novel object recognition test (NOR). p<0.05 by t-test comparing the novel vs. familiar object of each treatment/genotype. E) Time spent grooming in an open field. p < 0.05 for “genotype” by two-way ANOVA. For all tests, N=9-10 mice derived from 5 litters per condition.

### SCB alters PIN state by upregulating Shank 3 interactions

Next, we investigated whether SFK inhibition normalizes activity-dependent protein interaction networks in FMR1^-/y^ mice. To test the hypothesis that SCB would rescue the network pre-activation phenotype, we used cultured cortical neurons grown from FMR1^-/y^ and WT littermate controls, and pre-treated for 2 hours with SCB before stimulation with NMDA or DHPG (Fig 5A and Fig S8). In all groups, the average scaled value of the NMDA-responsive blue module was significantly reduced following NMDA treatment. Moreover, the module was significantly reduced in FMR1^-/y^ KO-aCSF compared to WT-aCSF, consistent with network pre-activation (Fig 5B). In FMR1^-/y^ KO-SCB-aCSF, the pre-activation was significantly greater than in KO FMR1^-/y^-aCSF, and less similar to the WT-aCSF cells (Fig 5B). A second module, brown, responded to DHPG in WT animals (Fig 5C), but did not respond to DHPG in FMR1^-/y^ with or without SCB treatment (Fig 5C). These data demonstrate that following acute 2-hour SCB treatment, network pre-activation is not normalized to WT levels in cultured cells, but rather becomes more divergent.

**Figure 5:**
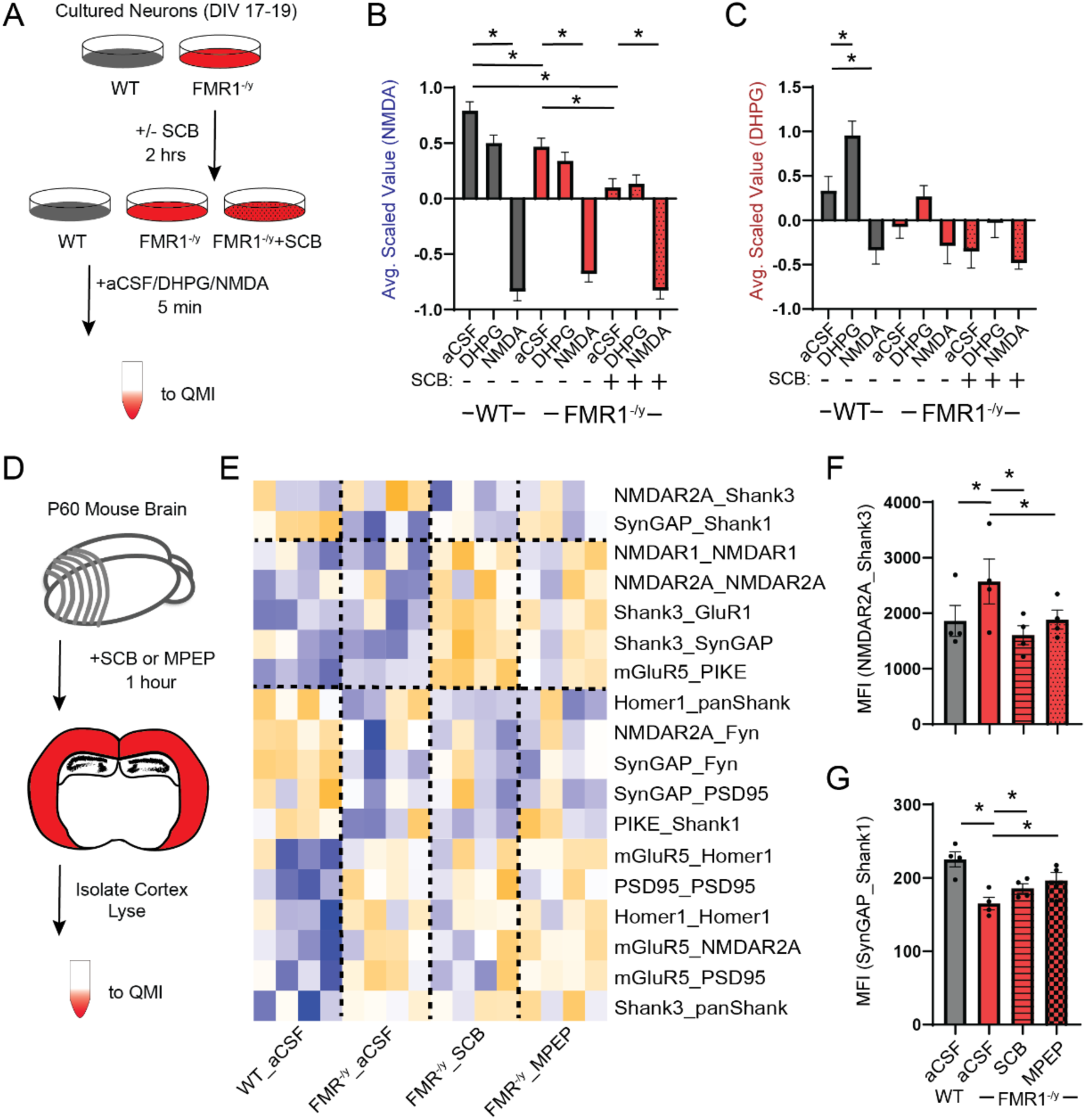
Acute SCB does not normalize protein interaction networks. **A)** Experimental design. B) The averaged scaled value of the NMDA-responsive blue module was significantly reduced following NMDA in all groups. FMR1^-/y^ showed significant module pre-activation, which was significantly exacerbated by SCB. C) The averaged scaled value of the DHPG-responsive brown module was significantly increased following DHPG in WT slices. In FMR1^-/y^ slices, the module did not respond to DHPG with or without SCB pretreatment. D) Experimental design. E) Interactions that were both individually statistically significant by a Bonferroni-corrected adaptive non-parametric test corrected for multiple comparisons (ANC) comparing treatment groups, and were a member of a CNA module that correlated with genotype, are represented by a row-scaled heatmap (blue-low abundance, orange-high abundance). Dotted lines divide interactions that were abnormal in FMR1^-/y^ slices and returned to WT levels by SCB and MPEP (top), those that were changed by SCB and MPEP (middle), and those that remained abnormal despite SCB/MPEP treatment (bottom). F,G) Two interactions were ‘normalized’ by SCB and MPEP treatment, including NMDAR2A_Shank3 (F) and SynGAP_Shank1 (G). MFI, median fluorescent intensity. Asterisks represents statistically significant difference by ANC, p<0.05. N = 4 WT and 4 FMR1^-/y^ animals; slices from each FMR1^-/y^ animal were treated with aCSF, SCB and MPEP to directly compare the effects of treatment.

Since signal transduction did not seem to be affected by SCB, we focused on protein interactions that may change in response to SCB treatment. Acute slices from P60 FMR1^-/y^ mice were treated with SCB for 1 hour, or with the mGluR5 antagonist MPEP to compare SCB treatment with another treatment that rescues protein synthesis and behavior in FMR1^-/y^ mice^52, 53^ (Fig 5D and Fig S8). Heatmapping of the single trait-associated module revealed three sets of interactions with distinct patterns of change: two interactions were significantly different between FMR1^-/y^ and WT aCSF slices, and were also significantly different between aCSF and drug-treated FMR1 slices, NMDAR2A_Shank3 and SynGAP_Shank1 (Fig 5 E-G). Several additional interactions involving NMDARs and Shank3 were not different between WT and FMR^-/y^ slices at baseline, but increased with MPEP and SCB treatment (e.g. NMDAR1_NMDAR1, Fig S8G). Finally, a number of interactions involving mGluR5, SynGAP and the Homer-Shank-PSD95 scaffold were up- or down-regulated in FMR1^-/y^ slices, and were not affected by drug treatment (Fig 5E). These latter interactions were overrepresented in the module such that the average scaled value of the module was significantly increased in FMR1^-/y^ animals, regardless of drug treatment. (Fig S8H). Importantly, MPEP and SCB induced a similar network of protein interaction alterations, suggesting that acute SCB treatment of FMR1^-/y^ mice acts on similar molecular pathway as MPEP treatment, possibly by decreasing Fyn activation downstream of hyperactive mGluR5 signaling. Overall, these data indicate that the gsPIN abnormalities observed in FMR1^-/y^ brain slices were not “normalized” to a WT-like state by acute treatment with SCB or MPEP.

To directly examine the effects of long-term SCB treatment *in vivo*, we dissected frontal cortex from mice orally gavaged with SCB for 7-14 days, or vehicle controls, and ran QMI (Fig 6A). A single trait-associated module correlated best with a model in which interactions were exacerbated, not rescued, by SCB treatment (Model 2, coded 0,1,2 for WT, FMR1^-/y^, FMR1^-/y^+SCB) (Fig 6B). Indeed, the average scaled value of the module increased from WT, to FMR1^-/y^, to FMR1^-/y^+SCB (Fig 6C). Heatmapping of the module (Fig 6D) revealed that the majority of interactions that were different between WT and FMR1^-/y^ mice, such as mGluR5_mGluR5 (Fig 6E) were not rescued by SCB treatment, and instead trended towards increasing differences from WT. Only two interactions were significantly upregulated in FMR1-/y mice, and significantly downregulated by SCB, Fyn_Fyn (likely reflecting direct inhibition by SCB, Fig 6F) and NMDAR2A_NL3. Thus, SCB treatment did not normalize protein interaction networks that were disrupted by FMR1 deletion.

**Figure 6:**
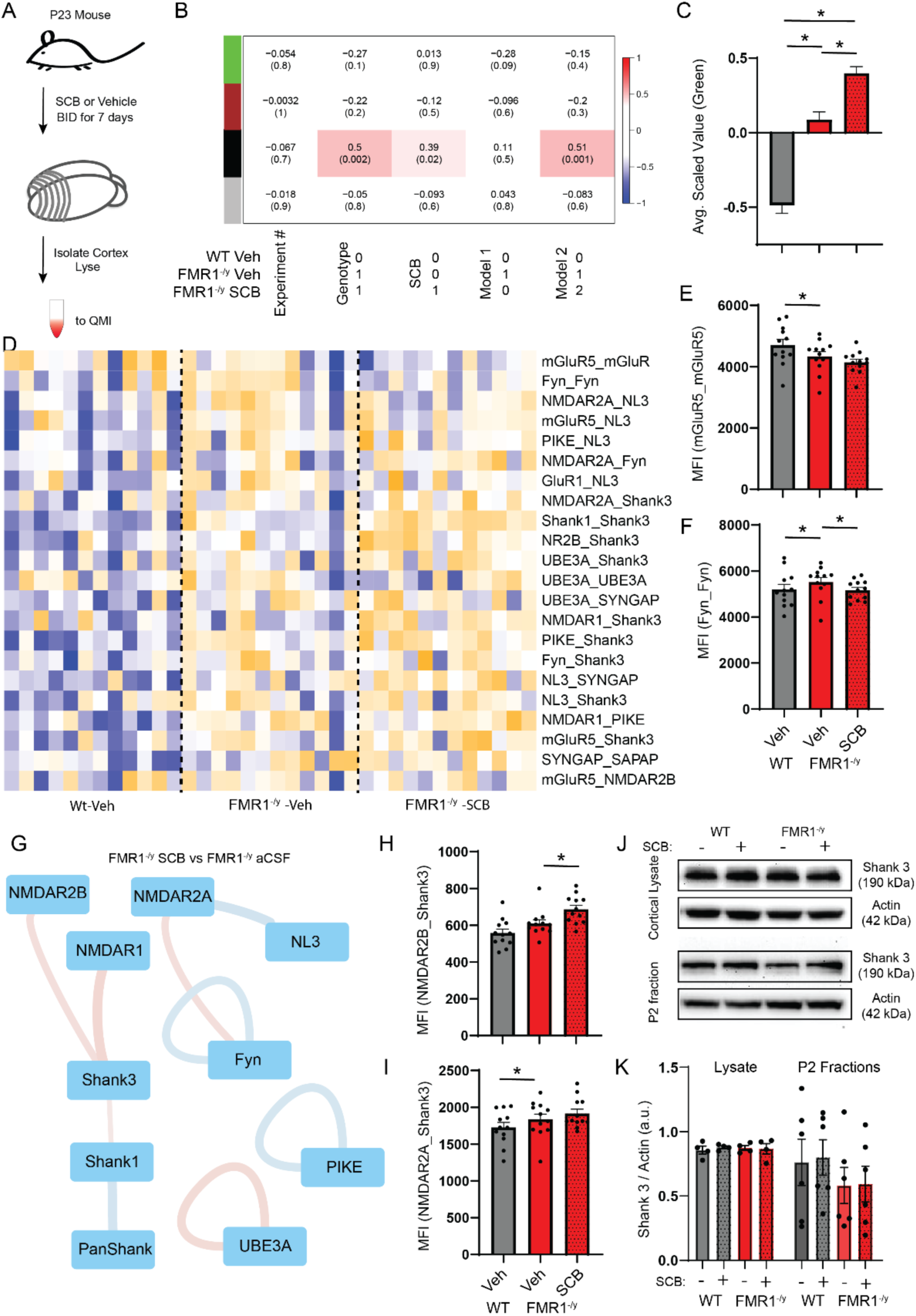
SCB does not normalize protein interaction networks *in vivo*. **A)** Experimental design. B) Module-trait correlation table shows the correlation coefficient (top number) and p-value (bottom number) of the correlation between each module eigenvector and the binary-coded hypothesis listed below the table. C) Averaged scaled value of all interactions in the genotype-and SCB-responsive black module. Asterisks represent statistically significant difference by one-way ANOVA followed by Sidak’s multiple comparison test, p<0.05. D) Interactions that were both individually statistically significant by a Bonferroni-corrected adaptive non-parametric test corrected for multiple comparisons (ANC) comparing treatment groups, and were a member of a significant module, are represented by a row-scaled heatmap (blue-low abundance, orange-high abundance). N = 12 animals per condition. E) mGluR5_mGluR5 was downregulated in FMR1^-/y^ animals, and not normalized by SCB. F) Fyn_Fyn was ‘normalized’ by SCB treatment. MFI, median fluorescent intensity. Asterisk represents statistically significant difference measured by ANC. G) Node-edge diagram shows protein interactions significantly different between untreated FMR1^-/y^ and FMR1^-/y^ SCB-treated slices. Nodes represent proteins, edges represent protein-protein interactions that were significantly different in the indicated comparison; red = increased, blue = decreased. The relative magnitude of the change is reflected in edge thickness. H,I) Shank3 interactions with NMDAR2B (H) and NMDAR2A (I) were increased in FMR1^-/y^ mice, and further increased by SCB treatment. Asterisks represents statistically significant difference by ANC, p<0.05. J) Representative Western blots and K) quantification of blots showing Shank3 expression in cortical lysates (N = 4 animals per condition) and P2 fractions (N = 6 animals per condition) from FMR1^-/y^ and WT mice treated with vehicle or SCB.

To determine potential mechanisms for the SCB behavioral rescue, we directly compared FMR1^-/y^ and FMR1^-/y^+SCB and found nine interactions that were significantly different, including interactions involving Shank3, NMDARs, and Fyn (Fig 6G). NMDAR2B_Shank3 was significantly increased (Fig 6H), and NMDAR2A_Shank3 was also elevated in FMR1^-/y^ mice (Fig 6I), as well as seven other interactions involving Shank3. Shank 3 is a scaffolding protein that plays an important role in organizing protein networks within the postsynaptic density of excitatory synapsis, and is critical for synaptic structure and function. However, western blot analysis revealed no changes in Shank 3 expression in cortical lysates or hippocampal P2 fractions from FMR1^-/y^ mice, with or without SCB (Fig 6J,K). Overall, these data demonstrate that acute or chronic SCB treatment, *in vitro* or *in vivo*, does not restore PINs to wildtype network states, but rather induces changes in protein interactions involving Fyn, Shank3 and NMDARs that correlate with improvements in protein synthesis and behavior.

## Discussion

### Modular organization of NMDA vs. DHPG signaling in FMR1^-/y^ animals

Rather than a static ‘scaffold’, the glutamate synapse is a liquid-liquid phase separated system^54, 55^ that, through phosphorylation or other post-translational modifications, alters the relative strength of its constitutive interactions. A property of liquid-liquid phase separated systems is that small changes in the relative binding affinity of system constituents can cause long-lasting changes in the size or membership of the overall system^54^. QMI allows sampling of the relative strength of hundreds of binary protein interactions, which provides a proxy measure of the overall gsPIN state. Our protein interaction network approach demonstrated that both WT and FMR1^-/y^ neurons and slices show changes in the relative abundance of modules of interactions following NMDA or DHPG stimulation. Both the NMDA- and DHPG-responsive modules showed evidence of pre-activation in FMR1^-/y^ neurons, in which the basal, unstimulated protein network state resembled the wildtype, neurotransmitter-stimulated state. In some cases, this pre-activation was so large that it occluded any further activation by DHPG, rendering FMR1^-/y^ neurons insensitive to DHPG. Comparing overall patterns of module activation in WT vs FMR1^-/y^ slices, this ‘pre-activation’ resulted in less contrast between the network states associated with aCSF, NMDA and DHPG conditions. We speculate that this reduction in the contrast and/or the dynamic range of the system may reduce its ability to distinguish between, and respond appropriately to, more physiologically relevant signaling inputs.

In prior studies of mature mouse hippocampal neurons from FMR1^-/y^ mice, excessive expression of Homer1a and excessive CAMKII-dependent phosphorylation of the Homer1 EVH1 domain^35, 56, 57^, lead to a dissociation of mGluR5 from the Shank-Homer scaffolding system which typically holds mGluR5 receptors in a peri-synaptic positon^58–60^. This disruption of mGluR5_Homer1 scaffolding allows the direct interaction of mGluR5 with NMDARs^61^, which serves to inhibit NMDA currents^57, 61, 62^. Consistent with a disrupted Homer-mGluR5 scaffold, in FMR1^-/y^ mice, mGluR5 is more abundant in the synaptic, rather than peri-synaptic, compartment^57^. Artificially localizing mGluR5 to the synapse in WT mice using an inducible protein interaction system increases spontaneous synaptic calcium transients^60^. An analogous increased synaptic calcium current in FMR1^-/y^ mice could be responsible for the ‘pre-activated’ state of the protein network, since mGluR5 signals through IP3 receptors to induce release of ER calcium stores. This high basal calcium activity could produce elevated CAMKII phosphorylation, phosphorylated Homer EVH1 domains, and activation of Homer1a translation, leading to a pathological feedback loop^16^ that maintains the gsPIN in a permanently altered state. Elevated calcium could also provide a mechanism for the activation of Fyn (and/or other SFKs) downstream of mGluR5.

### SFK inhibition as a treatment for Fragile X

In theory, the gsPIN could be reverted to a normal state if this pathological feedback loop could be disrupted. We focused on the kinase Fyn due the large number of dysregulated interactions containing Fyn, particularly at the critical developmental point of P17 when the DHPG module was clearly distinct from the NMDA module in WT animals. Fyn is a SRC-family tyrosine kinase that is widely expressed in the brain. Fyn was identified as a key kinase downstream of LTP at the same time as CAMKII^63^, but the field focused on CAMKII as a “learning and memory molecule” because its auto-phosphorylation mechanism led to the hypothesis that it could act as a molecular switch for memory^64^, while Fyn kinase was largely relegated to the background. However, Fyn is not only critical for LTP^65^, but also for spatial learning^63^, contextual fear conditioning^66^, and alcohol self-administration^67^. Fyn knockout mice show decreased dendritic spine density and altered spine morphology^68^, abnormal spatial learning, and impaired LTP^63^ (but see^69^). Fyn physically interacts with type 1 mGluRs^38, 70^, and phosphorylates both NMDA^40^ and AMPA^71^ receptors following synaptic activity. Thus, Fyn is a central, and arguably understudied, regulator of synaptic plasticity that is disrupted in FMR1^-/y^ mice.

We selected SCB to inhibit Fyn due to its promising pharmacological profile, having been safely administered to hundreds of study participants in trials for Cancer^72, 73^ and Alzheimer’s disease^49^. While SCB is not an FDA approved drug because those trails did not show clinical efficacy, the side effects were generally mild and well tolerated. Moreover, robust mouse studies have identified dosing strategies that produce CSF levels in mice that are similar to steady-state levels in human trails^42^, making our dosing strategy physiologically relevant. Due to the high degree of homology among SRC family kinases, particularly in the kinase domain itself, there are no available drugs that inhibit Fyn and do not cross-react with other SFKs. Similarly, there are no antibodies for phospho-Fyn, so we had to rely on phospho-SFK. However, we took the approach of using a drug that targeted Fyn as well as other SFKs, but that has a well-established safety and dosing profile, with the goal of facilitating translational, rather than purely mechanistic, studies. Follow-up studies using genetic approaches should be conducted to isolate the effect of Fyn as opposed to the other brain-expressed SRC family kinases such as Src, Lyn, or Yes.

Many different treatments have been shown to improve behavior in the FMR1^-/y^ mouse: Inhibition of mGluR5^52, 74, 75^, Ras-Erk^21, 76–78^ signaling (but not mTOR^79^) and GSK3a^80^ all produce acute rescue of protein synthesis and behavioral phenotypes. Further, CAMKII inhibition improves seizures^35^, and knockdown of Homer1a by AAV improves memory in FMR1^-/y^ mice^57^. All of these phenotypes could lie downstream of hyperactivation of the gsPIN, and our comparison of MPEP and Fyn treatment in acute FMR1^-/y^ slices suggests that Fyn and MPEP share similar mechanisms. In addition to identifying a novel potential drug to treat Fragile X symptoms, our findings that SCB rescued excessive protein synthesis *in vitro* and rescued select behavioral deficits *in vivo* are important because they act as a proof-of-principle that analysis of the protein interaction network downstream of signaling inputs can identify unforeseen drug targets.

### Behavioral normalization does not correlate with gsPIN normalization

Our initial, intuitively obvious hypothesis was that if abnormal protein interaction networks correlate with abnormal molecular and behavioral phenotypes, then treatments that normalize these downstream phenotypes should also normalize the protein network status. We tested this hypothesis with both SCB and MPEP, and in both cases found that it was false. Instead, the gsPIN state appeared, by many network measures, more abnormal than the untreated FMR1^-/y^ state. Very few interactions met the criteria of being significantly different than WT in the untreated mutant, and normalized by SCB or MPEP treatment. Rather, we identified a network of interactions involving Fyn, Shank3 and NMDARs that were altered by SCB, and that might be associated with molecular and behavioral rescue. Since we showed that SCB treatment does not change Shank3 expression in WT and FMR1^-/y^ mice, we speculate that SCB may affect Shank3 phosphorylation profiles. Given that phosphorylation of scaffolding proteins is an important regulatory mechanism for protein interactions, subcellular distribution, and physiological functions, and that Shank3 is a highly phospho-regulated protein^81, 82^, further studies will be necessary to unravel how SCB treatment affects Shank 3 phosphorylation state and protein interactions.

## Materials and Methods

### Animals

Male FMR1 knockout (FMR1^-/y^) and wildtype (WT) littermates on the C57BL/6J background used studied in all experiments. The breeding scheme was female FMR1 heterozygous mice (JAX #003025) crossed with wildtype male C57BL/6J mice (JAX#000664). Age-matched littermates were randomized to treatment groups and a balanced number of FMR1^-/y^ and WT mice were used. Mice were group-housed with no more than 5 mice/cage and maintained on a 12:12 h light/dark cycle. Food and water were provided *ad libitum*. All work was performed under an approved protocol by the Office of Animal Care at the Seattle Children’s Research Institute (protocol #00072).

### Cell Culture

Primary cultures of cortical neurons were prepared as previously described^5, 6^. Briefly, whole cortex from P0-P1 mouse neonate was dissociated using papain (Worthington Biochemical) and plated at a density of 1-1.2x10^6^ cells/ml onto 6-well plates treated with poly-D-lysine (EMD Millipore). Cells were cultured in Neurobasal A medium supplemented with 2% B27, 2mM GlutaMax, 100U/ml penicillin, and 100µg/ml streptomycin (ThermoFisher) and kept at 37°C, 5% CO_2_ for 18-21 days. After 3-5 DIV, 5-fluoro-2’-deoxyuridine was added to a final concentration of 5 µM to inhibit glial proliferation.

### Neuronal stimulation and lysate preparation

For direct activation of glutamate receptors, neurons were treated with NMDA/glycine (100/1 µM), DHPG (100 µM) or control HEPES-aCSF (129 mM NaCl/ 5 mM KCl/ 2 mM CaCl_2_/ 30 mM Glucose/ 25 mM HEPES; pH 7.4) for 5 minutes. NMDAR agonists NMDA (Sigma, M3262; 100 µM) + glycine (Fisher Scientific, Pittsburgh, PA, USA, BP-381; 1 µM) were dissolved in HEPES-aCSF in the presence of non-NMDA glutamate receptor blockers (CNQX, Tocris, 0190, 40 µM; Nimodipine, Tocris, 0600, 1 µM; LY-367385, Sigma, L4420, 100 µM; MPEP, Tocris, 1212, 10 µM). Type I mGluR agonist DHPG (Tocris, 0805; 100 µM) was dissolved in HEPES-aCSF in the presence of non-mGluR glutamate receptor blockers (CNQX, Tocris, 0190, 40 µM; Nimodipine, Tocris, 0600, 1 µM; D(-)-2-Amino-5 phosphonopentanoic acid (APV), Sigma, A5282, 50 µM). For testing the effect of SCB, neurons were first pre-treated with 2µM SCB (dissolved in DMSO) for 2 hrs, and the same concentration of the drug was added in the stimulation buffers. Following treatment, cells were immediately placed on ice and rapidly harvested by scraping in lysis buffer (150mM NaCl, 50mM Tris (pH 7.4), 1% NP-40, 10 mM sodium fluoride (Sigma, 201154), 2 mM sodium orthovanadate (Sigma, 450243) + protease/phosphatase inhibitor cocktails (Sigma, P8340/P5726)), followed by incubation in lysis buffer for 15 minutes. Lysate was centrifuged for 15 minutes at high speed to remove nuclei and debris, and protein concentration in the supernatant was determined using a Pierce BCA kit (ThermoFisher, 23225).

### Slice preparation, stimulation, and lysate preparation

Acute brain slices were prepared from P7, P17 and P60 FMR1^-/y^ and WT littermate mice using N-methyl-D-glucamine (NMDG) protective recovery method ^83^. Mice were deeply anesthetized with isoflurane, brains were removed, and 400 μm-thick coronal cortical slices were sectioned in pre-chilled carbogenated NMDG-HEPES aCSF (92 mM NMDG, 2.5 mM KCl, 1.2 mM NaH_2_PO_4_, 30 mM NaHCO_3_, 20 mM HEPES, 25 mM glucose, 2 mM thiourea, 5 mM Na-ascorbate, 3 mM Na-pyruvate, 0.5 mM CaCl_2_.4H_2_O, and 10 mM MgSO_4_.7H_2_O; titrated to pH 7.4 with concentrated hydrochloric acid) using a VT1200s vibrating microtome (Leica). The transcardial perfusion procedure with 30 ml of chilled NMDG-HEPES aCSF was performed on P60 animals prior brain dissection and slicing. Slices were immediately hemisected with a sharp razor blade and each half placed in an alternate treatment group, with treatment groups being arbitrarily assigned. Upon completion of the sectioning procedure, all of the slices were collected using a cut-off plastic Pasteur pipet and transferred into a pre-warmed (32-34°C) initial recovery chamber filled with NMDG-HEPES aCSF and incubated for 10-15 min. Slices were then transferred to a carbogenated modified HEPES holding solution (92 mM NaCl, 2.5 mM KCl, 1.2 mM NaH_2_PO_4_, 30 mM NaHCO_3_, 20 mM HEPES, 25 mM glucose, 2 mM thiourea, 5 mM Na-ascorbate, 3 mM Na-pyruvate, 2 mM CaCl_2_.4H_2_O, and 2 mM MgSO_4_.7H_2_O; pH 7.4) for an additional 60–90 min recovery at room temperature. Slices were then incubated in HEPES-aCSF (129 mM NaCl/ 5 mM KCl/ 2 mM CaCl_2_/ 30 mM Glucose/ 25 mM HEPES; pH 7.4) for 1h at 32–34 °C to equilibrate, and subsequently stimulated with NMDA/glycine (100/1 μM), DHPG (100 uM) or HEPES-aCSF control for 5 minutes at 32–34°C. Similar to cell culture experiment, antagonists for non-targeted receptors were added in the stimulation buffer to isolate the targeted receptor. For testing the effects of SCB and MPEP, the same slice recovery procedure was applied, after which 2 µM SCB (Cayman Chemical, 11497) or MPEP (Tocris, 1212) was added in HEPES-aCSF and incubated for 1h at 32–34 °C. In this experiment only adult mice were used and no glutamate receptor stimulation was performed. Following treatments, slices were quickly frozen in liquid nitrogen and kept at -80 C until further processing. Lysates from cortical slices were prepared as above with the following exception: tissue was homogenized in ice-cold 1% NP-40 lysis buffer using 12 strokes of a glass-teflon homogenizer (PYREX).

### Quantitative Multiplex co-Immunoprecipitation (QMI)

QMI was performed as described previously^5–7^, see Brown et al^33^ for a detailed video protocol, analysis software code for both R and Matlab, and sample datasets. QMI validation in mouse neurons, including specific validation of each antibody pair, was described previously^5^. Antibody clone names and catalog numbers are the same as in^7^, and are listed in Table S1.

To perform QMI, a master mix containing equal numbers of each antibody-coupled Luminex bead was prepared, distributed to lysates containing equal amounts of protein, and incubated overnight on a rotator at 4°C. The next day, beads from each sample were washed twice in cold Fly-P buffer (50 mM tris (pH7.4), 100 mM NaCl, 1% bovine serum albumin, and 0.02% sodium azide) and distributed into twice as many wells of a 96-well plate as there were probe antibodies (for technical duplicates). Biotinylated detection (probe) antibodies were added to the wells and incubated at 4°C with gentle agitation for 1 hour. The resulting bead-probe complexes were washed three times with Fly-P buffer, incubated for 30 min with streptavidin-phycoerythrin on ice, washed another three times, resuspended in 125 μl of ice-cold Fly-P buffer, and read on a customized refrigerated Bio-Plex 200 (as detailed in ref^33^).

### Quantification and statistical analysis of QMI data

#### Data preprocessing and inclusion criteria

For each well from a data acquisition plate, data were processed using the BioPlex200’s built-in software to (i) eliminate doublets on the basis of the doublet discriminator intensity (> 5000 and < 25 000 arbitrary units; Bio-Plex 200), (ii) identify specific bead classes within the bead regions used, and (iii) pair individual bead phycoerythrin fluorescence measurements with their corresponding bead regions. This processing generated a distribution of fluorescence intensity values for each pairwise protein interaction measurement. XML output files were parsed to acquire the raw data for use in MATLAB using the ANC program^36^. No specific analysis was performed on the data to test for outliers.

#### Adaptive non-parametric analysis with empirical alpha cutoff (ANC)

ANC is used to identify high-confidence, statistically significant differences (corrected for multiple comparisons) in bead distributions for each pairwise protein interaction. ANC analysis was conducted in MATLAB (version 2013a) as described in^36^. As previously reported, we required that hits must be present in > 70% of experiments (typically three out of four) at an adjusted p < 0.05. The a-cutoff value required per experiment to determine statistical significance was calculated to maintain an overall type I error of 0.05 (adjusted for multiple comparisons with Bonferroni correction), with further empirical adjustments to account for technical variation. No assessment of normality was carried out as ANC analysis is a non-parametric test.

#### Correlation Network Analysis (CNA)

Modules of protein-protein interactions that co-varied with experimental conditions were identified using CNA as described in ^33, 36^. Briefly, bead distributions used in ANC were collapsed into a single median fluorescent intensity (MFI) for each interaction and averaged across technical replicates for input into the WGCNA package for R^34^. Interactions with MFI < 100 were removed as noise, and batch effects were corrected using COMBAT^84^ with ‘experiment number’ as the batch. Power values giving the approximation of scale-free topology were determined using soft thresholding with a power adjacency function. The minimum module size was set to 10, and modules whose eigenvectors significantly correlated with an experimental trait (p < 0.05) were considered ‘‘of interest.’’ Interactions belonging to a module of interest and whose probability of module membership in that module was < 0.05 were considered significantly correlated with that trait.

#### Principal Component Analysis (PCA)

PCA was performed on Post-COMBAT, log2 transformed MFI values in R studio using the prcomp function.

#### ANCՈCNA

Interactions that were significant by both ANC and CNA for a given experimental condition were considered significantly altered in that condition. Only these ‘double-significant’ interactions are reported in the text and figures.

#### Data visualization

For heatmap visualization of MFI values of CNA module members, log2 transformed data were input into the Heatmap.2 program in R studio, which normalized the data by row for visualization of multiple analytes spanning a 3-log range. All interactions visualized were ANCՈCNA significant for at least one condition. The number of biological replicates for each experiment can be found in figure legends, error bars represent SEM.

### Puromycin assay

Basal protein synthesis rates were measured in acute cortical slices from P21-23 FMR1^-/y^ and WT littermate mice. Mice were deeply anesthetized with isoflurane, decapitated and brains were rapidly removed. Three 400 µm-thick prefrontal cortical slices were cut in ice-cold aCSF (129 mM NaCl/ 5 mM KCl/ 2 mM CaCl_2_/ 1 mM MgCl_2_/ 30 mM Glucose/ 25 mM HEPES; pH 7.4) with 95% O_2_-5% CO_2_ using a VT1200s vibrating microtome (Leica). Following cutting, each slice was divided into two hemisphere and left for recovery period of 3.5 h in aCSF at 32-34°C (saturated with 95% O_2_-5% CO_2_) and then incubated in actinomycin D (1 µg/ml in DMSO) for 30 min at 32-34°C to inhibit transcription. Control slices were then incubated in solution containing actinomycin D (1 µg/ml) and puromycin (5 µg/ml) for 1 h at 32-34°C, whereas for SCB-treated slices 2 µM SCB (Cayman Chemical, 11497) was added in actinomycin D/puromycin incubation solution and incubated for the same incubation period. Slices were washed 3 times in ice cold aCSF to rinse of excess puromycin, and then homogenized with 12 strokes of a glass-Teflon homogenizer with twisting in 1% deoxycholate lysis buffer with 0.1M Tris-HCl (pH 9), 10 mM sodium fluoride (Sigma, 201154), 2mM sodium orthovanadate (Sigma, 450243) and protease/phosphatase inhibitor cocktails (Sigma, P8340/P5726). Following incubation in lysis buffer for 15 min, lysates were centrifuged for 15 min at 13,000 G to remove nuclei and debris. Protein concentration in the supernatant was determined using a Pierce BCA kit (ThermoFisher, 23225). Samples were analyzed by immunoblot, and puromycin incorporation was detected using the mouse monoclonal anti-puromycin (12D10) antibody (1:1000, Millipore, MABE343). In addition, PVDF membranes were probed for rabbit anti-Src family (Y416) antibody (1:1000, Cell Signaling, 2101), rabbit anti-Fyn (1:1000, Cell Signaling, 4023), rabbit anti-alpha FMR1 (c-terminal) antibody (1:2000, Sigma, F4055) and rabbit-anti GAPDH antibody (1:10,000, Cell Signaling, 2118). Primary antibodies were detected using species-specific horseradish peroxidase–conjugated secondary antibodies.

### Behavioral testing

#### Drug treatment

A large cohort (N∼40) of P23-26 FMR1^-/y^ mice, or their WT littermates, were treated with saracatinib (SCB, 5 mg/kg/d, Cayman Chemical, 11497) or vehicle control (Veh; 0.5% w/v HPMC/0.1% w/v Polysorbate 80 in saline) by oral gavage for 7 days prior to the start of behavioral testing and the treatment was continued throughout the duration of the behavioral testing. The dose and method of drug administration were as described in^42^. Since the estimated half-life of SCB in mice is 16 hours, the daily dose was divided in two equal amounts. Oral delivery at 5 mg/kg/d of SCB yields levels expected to inhibit at least 50% of Fyn kinase activity in the brain^42^. All behavioral testing was done blind to genotypes and treatments during the light cycle (12 - 6 pm).

#### Open field test

Each mouse was gently placed in a corner of a 60 cm x 60 cm open-field arena illuminated with white light, and the movement of the mouse was recorded by a ceiling-mounted camera for 10 min. The recorded video file was further analyzed using EthoVision XT 7.0 software (Noldus). Total distance moved, mean velocity, the number of entries into and the overall time spent in the center of the arena (40 × 40 cm imaginary square), as well as time spent grooming were measured. The open field arena was cleaned with 70% ethanol between each trial.

#### Novel object recognition test

A mouse was first habituated to an arena (30 cm x 30 cm) for 5 min. Following habituation, the mouse was removed from the arena and two identical objects were placed in the opposite corners of the arena, 7 cm from the side walls. The mouse was reintroduced into the center of the arena and allowed to explore the arena including the two novel objects for 10 min. After the retention period of 48 h, one object was replaced with another novel object, which was of similar size but different shape and color than the previous object. The mouse was placed in the arena and allowed to explore the two objects for additional 10 min. The movement of mice was recorded by a camera and the time spent interacting with the novel vs. familiar object was analyzed by the video tracking EthoVision XT 7 software (Noldus). The arena was cleaned with 70% ethanol between trials.

#### Three-chamber sociability test

Social behavior was evaluated by means of the three-chamber apparatus. A rectangular and transparent Plexiglas box (40 cm x 60 cm) divided by walls into three equal-sized compartments was used. The test mouse was placed in the center chamber of a three-chamber apparatus and allowed to explore all three chambers for a 5 min habituation period. Following habituation, an empty holding enclosure was introduced to one of the side chambers and another identical enclosure with a stranger mouse inside was introduced to the other side. The test mouse was allowed to explore the entire apparatus for another 5 min. Time spent in each chamber and time spent sniffing the stranger mouse enclosure were recorded. All apparatus chambers were cleaned with 70% ethanol between trials.

### P2 fractionation

Mice were deeply anesthetized with isoflurane, brains were removed, and 3 mm of frontal cortical tissue was cut with a razor blade. Tissue was frozen in liquid nitrogen and stored at -80 until homogenization. Cortices were then homogenized in 0.32M sucrose in 4 mM 4-(2-hydroxyethyl)-1-piperazineethanesulfonic acid (HEPES) buffer with protease/phosphatase inhibitor cocktails using 12 strokes of a glass-tefon homogenizer. The homogenate was immediately centrifuged at 1,000G for 10 min at 4°C. The supernatant (S1) was then spun at 12,000G for 15 min at 4 °C. The resulting P2 pellet was then resuspended in 1% deoxycholate lysis buffer for 15 min. Lysate was then centrifuged at 12,000G for 15 min to remove insoluble portions and protein concentration in the supernatant was determined using a Pierce BCA kit (ThermoFisher, 23225).

### Western Blotting

For Western blots, proteins (20 μg per lane) were separated by SDS–polyacrylamide gel electrophoresis and transferred to a polyvinylidene difluoride membrane (Bio-Rad). Membranes were blocked in 4% milk in TBST [0.01 M tris (pH7.4), 0.15 M NaCl, 0.1% Tween 20] for 1 hour at room temperature and incubated with primary antibodies overnight at 4°C. Primary antibodies and dilutions used for Western blots were Shank 3 (1:1000, clone N367/62, Neuromab), and β-actin (1:10,000, GTX109639, GeneTex). Primary antibodies were detected using species-specific horseradish peroxidase– conjugated secondary antibodies. Blots were developed using SuperSignal West Pico PLUS Substrate (ThermoFisher) and imaged using a ProteinSimple imaging system (San Jose, CA). Protein bands were measured by densitometry and quantified as the densitometric signal of Shank 3 divided by the densitometric signal of β-actin as loading control.

### Statistical analysis

Statistical test used in each experiment is denoted in the figure legends. All the statistical tests and graphs were performed using GraphPad Prism 9 (GraphPad Software Inc). For analysis of QMI data using statistical approaches ANC and CNA, see above. All data are expressed as mean ± SEM. N values representing numbers of animals are stated in the figure legends.

## Acknowledgements

This work was supported by NIH grants MH113545 and MH121487 (to SEPS), a fellowship from the FRAXA research foundation (to VS), and a NARSAD grant from the Brain and Behavior Research Foundation (to SEPS). The authors thank all members of the SEPS laboratory, as well as Drs. B.K. Reed and M. Chau for providing insight and thoughtful discussions.

## Author Contributions

Conceptualization: SEPS and JDL. Investigation: VS, JDL, FH. Data analysis: VS, JL, SEPS. Writing: VL and SEPS. Funding acquisition and supervision: VS and SEPS.

