## Supplemental Figures and Tables for "SRC family kinase inhibition rescues molecular and behavioral phenotypes, but not protein interaction network dynamics, in a mouse model of Fragile X syndrome"

**for**

**Includes:**

**Supplemental Figures S1-S8**

**Supplemental Table 1**

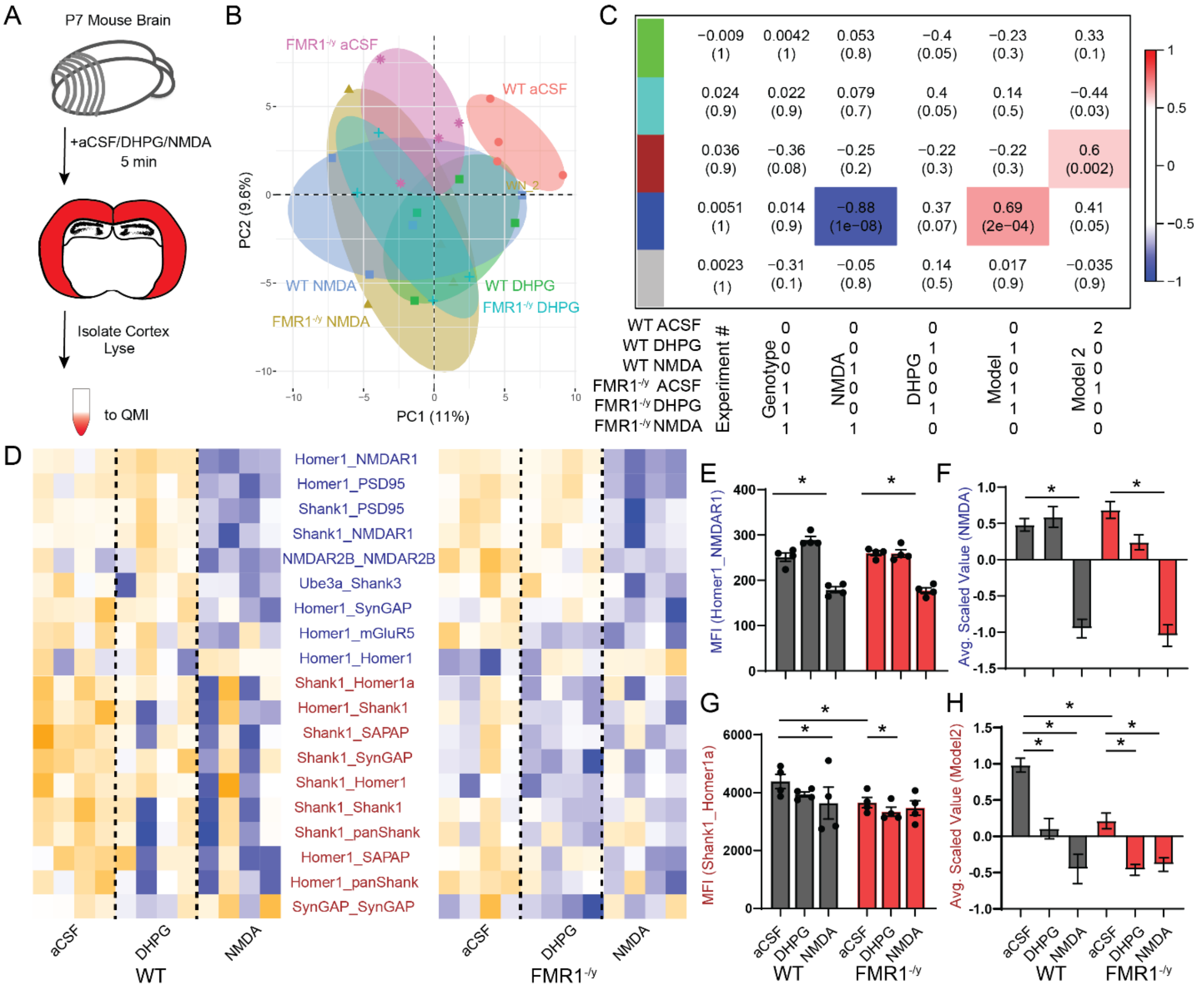


**Figure S1: FMR1$^{-/y}$ acute P7 brain slices show hyperactivation of DHPG-responsive protein interactions.** A) Experimental design. B) Principal component graph showing separation of wildtype aCSF (red), from NMDA (blue) and DHPG (green) treatment groups across PC1 and PC2. In FMR1$^{-/y}$ slices, aCSF treatment (magenta) is separate from WT_aCSF treatment, suggesting tonic activation. C) Module-trait correlation table shows the correlation coefficient (top number) and p-value (bottom number) of the correlation between each module eigenvector and the binary-coded hypothesis listed below the table. The blue module correlates with NMDA treatment and with a hypothesis of basal DHPG hyperactivation in FMR1$^{-/y}$ animals, while the brown module correlates with a model of partial activation in FMR1$^{-/y}$ - aCSF of dissociations that occur in response to NMDA and DHPG. D) Interactions that were both individually statistically significant by a Bonferroni-corrected adaptive non-parametric test corrected for multiple comparisons (ANC, see methods) comparing treatment groups, and were a member of a significant module, are represented by a row-scaled heatmap (blue- low abundance, orange- high abundance). Interactions are colored based on their assigned module. E) Example of an interaction in the NMDA-responsive blue module, Homer1_NMDAR1. MFI, median fluorescent intensity. Asterisks represent statistically significant difference measured by ANC. F) Averaged scaled value of all interactions in the NMDA-responsive blue module. Asterisks represent statistically significant difference by one-way ANOVA followed by Sidak's multiple comparison test, $p<0.05$. G) Example of an interaction in the DHPG-responsive brown module, Shank1_Homer1a. Asterisks represent statistically significant difference by ANC, $p<0.05$. H) Averaged scaled value of all interactions in the DHPG-responsive brown module. Asterisks represent statistically significant difference by one-way ANOVA followed by Sidak's multiple comparison test, $p<0.05$. N = 4 mice per condition

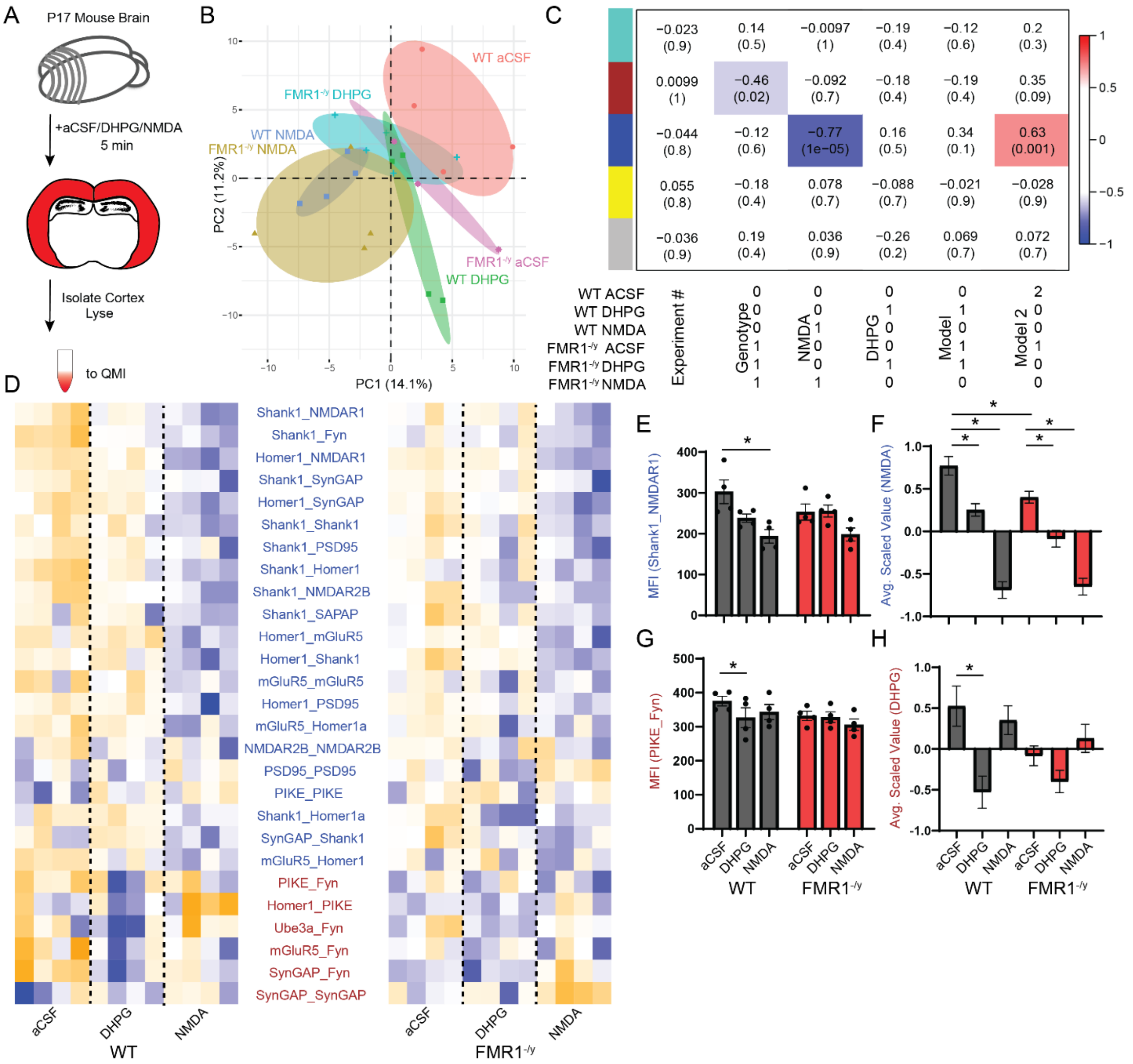

**Figure S2: FMR1$^{-/y}$ acute P17 brain slices show hyperactivation of DHPG-responsive protein interactions.** A) Experimental design. B) Principal component graph showing separation of wildtype aCSF (red), from NMDA (blue) and DHPG (green) treatment groups across PC1 and PC2. In FMR1$^{-/y}$ slices, aCSF treatment (magenta) overlaps with both WT_DHPG and FMR1$^{-/y}$_DHPG slices, suggesting tonic activation. C) Module-trait correlation table shows the correlation coefficient (top number) and p-value (bottom number) of the correlation between each module eigenvector and the binary-coded hypothesis listed below the table. The blue module correlates with NMDA treatment and with a hypothesis of partial activation in FMR1$^{-/y}$_aCSF of dissociations that occur in response to NMDA and DHPG. The brown module correlates with genotype. D) Interactions that were both individually statistically significant by a Bonferroni-corrected adaptive non-parametric test corrected for multiple comparisons (ANC, see methods) comparing treatment groups, and were a member of a significant module, are represented by a row-scaled heatmap (blue- low abundance, orange- high abundance). Interactions are colored based on their assigned module. E) Example of an interaction in the NMDA-responsive blue module, Shank1_NMDAR1. MFI, median fluorescent intensity. Asterisk represents statistically significant difference measured by ANC. F) Averaged scaled value of all interactions in the NMDA-responsive blue module. Asterisks represent statistically significant difference by one-way ANOVA followed by Sidak's multiple comparison test, $p<0.05$. G) Example of an interaction in the DHPG-responsive brown module, PIKE_Fyn. Asterisk represents statistically significant difference by ANC, $p<0.05$. H) Averaged scaled value of all interactions in the DHPG-responsive brown module. Asterisks represent statistically significant difference by one-way ANOVA followed by Sidak's multiple comparison test, $p<0.05$. N = 4 mice per condition.

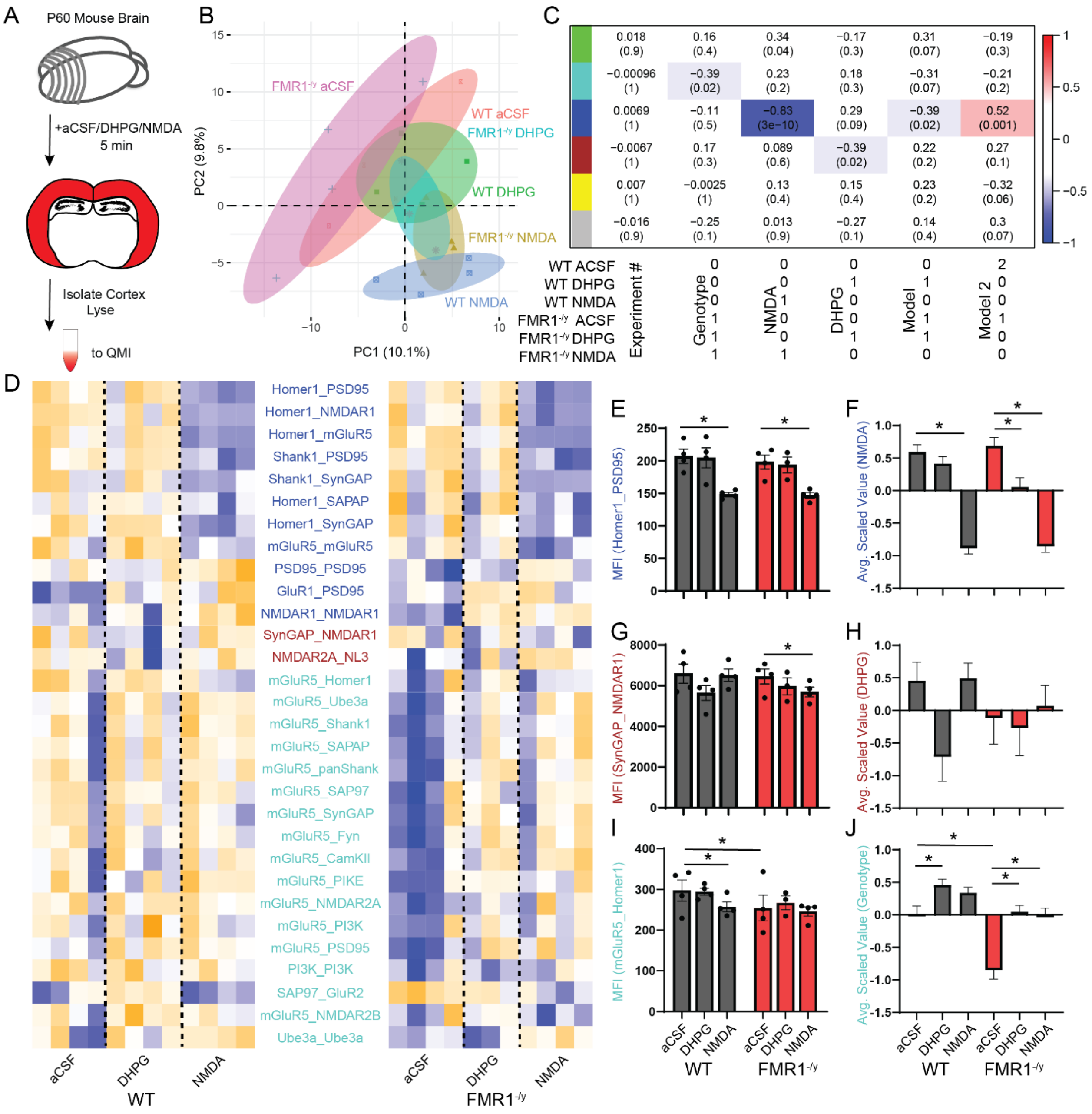


**Figure S3: FMR1$^{-/y}$ acute P60 brain slices show hyperactivation of DHPG-responsive protein interactions.** A) Experimental design. B) Principal component graph showing wildtype aCSF (red), NMDA (blue) and DHPG (green) treatment groups separating across PC1 and PC2. FMR1$^{-/y}$ slices occupied similar PCA space. C) Module-trait correlation table shows the correlation coefficient (top number) and p-value (bottom number) of the correlation between each module eigenvector and the binary-coded hypothesis listed below the table. The blue module correlates with NMDA treatment and with a hypothesis of partial activation in FMR1$^{-/y}$_aCSF of the dissociations that occur in response to NMDA and DHPG. The brown module correlates with DHPG treatment and turquoise with genotype. D) Interactions that were both individually statistically significant by a Bonferroni-corrected adaptive non-parametric test corrected for multiple comparisons (ANC, see methods) comparing treatment groups, and were a member of a significant module, are represented by a row-scaled heatmap (blue- low abundance, orange- high abundance). Interactions are colored based on their assigned module. E) Example of an interaction in the NMDA-responsive blue module, Homer1_PSD95. MFI, median fluorescent intensity. Asterisk represents statistically significant difference measured by ANC. F) Averaged scaled value of all interactions in the NMDA-responsive blue module. Asterisks represent statistically significant difference by one-way ANOVA followed by Sidak's multiple comparison test, $p<0.05$. G) Example of an interaction in the DHPG-responsive brown module, SynGAP_NMDAR1. Asterisk represents statistically significant difference by ANC, $p<0.05$. H) Averaged scaled value of all interactions in the DHPG-responsive brown module. I) Example of an interaction in the genotype-correlated turquoise module, mGluR5_Homer1. Asterisk represents statistically significant difference by ANC, $p<0.05$. J) Averaged scaled value of all interactions in the genotype-correlated turquoise module. Asterisks represent statistically significant difference by one-way ANOVA followed by Sidak's multiple comparison test, $p<0.05$. N = 4 mice per condition (3 for $FMR1^{-/y}$_DHPG due to an outlier being eliminated).

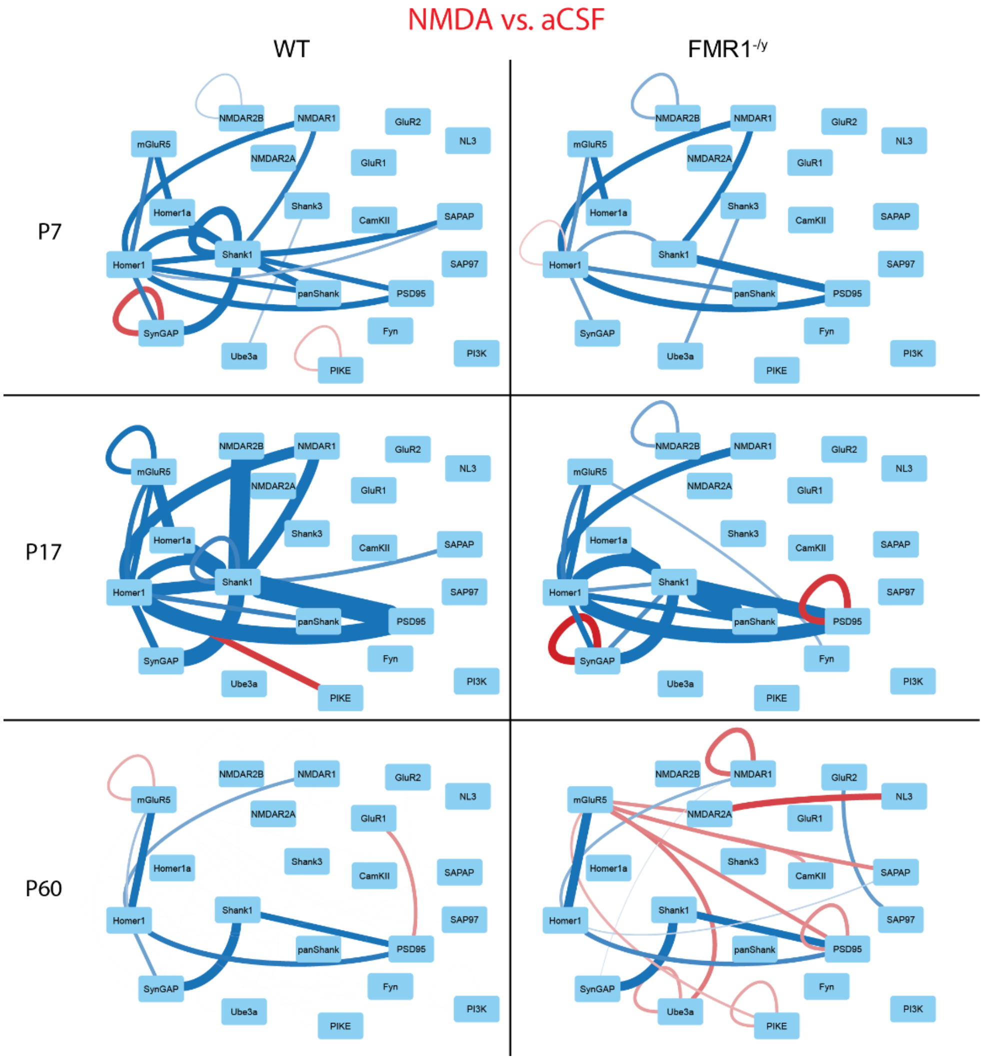


**Figure S4: NMDA-responsive protein interactions**. Node-edge diagrams show protein interactions that changed in response to NMDA stimulation (compared to aCSF) in WT or FMR1$^{-/y}$ slices from P7, P17 or P60 mice. Nodes represent proteins, edges represent protein-protein interactions that changed in response to NMDA; red = increased, blue = decreased. The relative magnitude of the change is reflected in edge thickness. All interactions shown were both statistically significant by a Bonferroni-corrected adaptive non-parametric test corrected for multiple comparisons (ANC), and were a member of a significant module. N = 4 animals per age/condition.

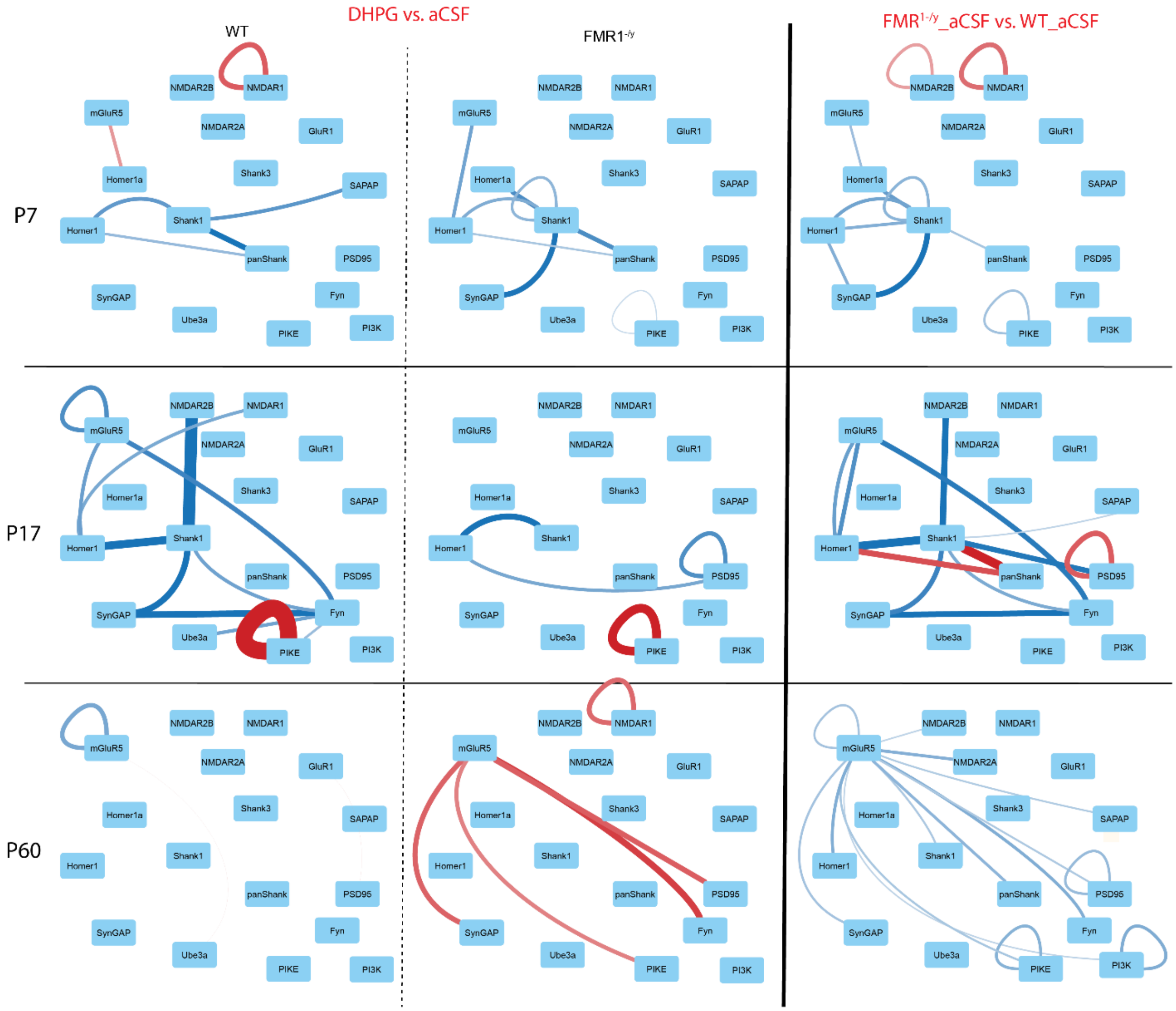


**Figure S5: DHPG-responsive protein interactions**. Node-edge diagrams show protein interactions that changed in response to DHPG stimulation (compared to aCSF), or that were different in comparisons of WT_aCSF and FMR1$^{-/y}$_aCSF, from P7, P17 or P60 mice. Nodes represent proteins, edges represent protein-protein interactions that changed in response to NMDA; red = increased, blue = decreased. The relative magnitude of the change is reflected in edge thickness. All interactions shown were both statistically significant by a Bonferroni-corrected adaptive non-parametric test corrected for multiple comparisons (ANC), and were a member of a significant module. N = 4 animals per age/condition.

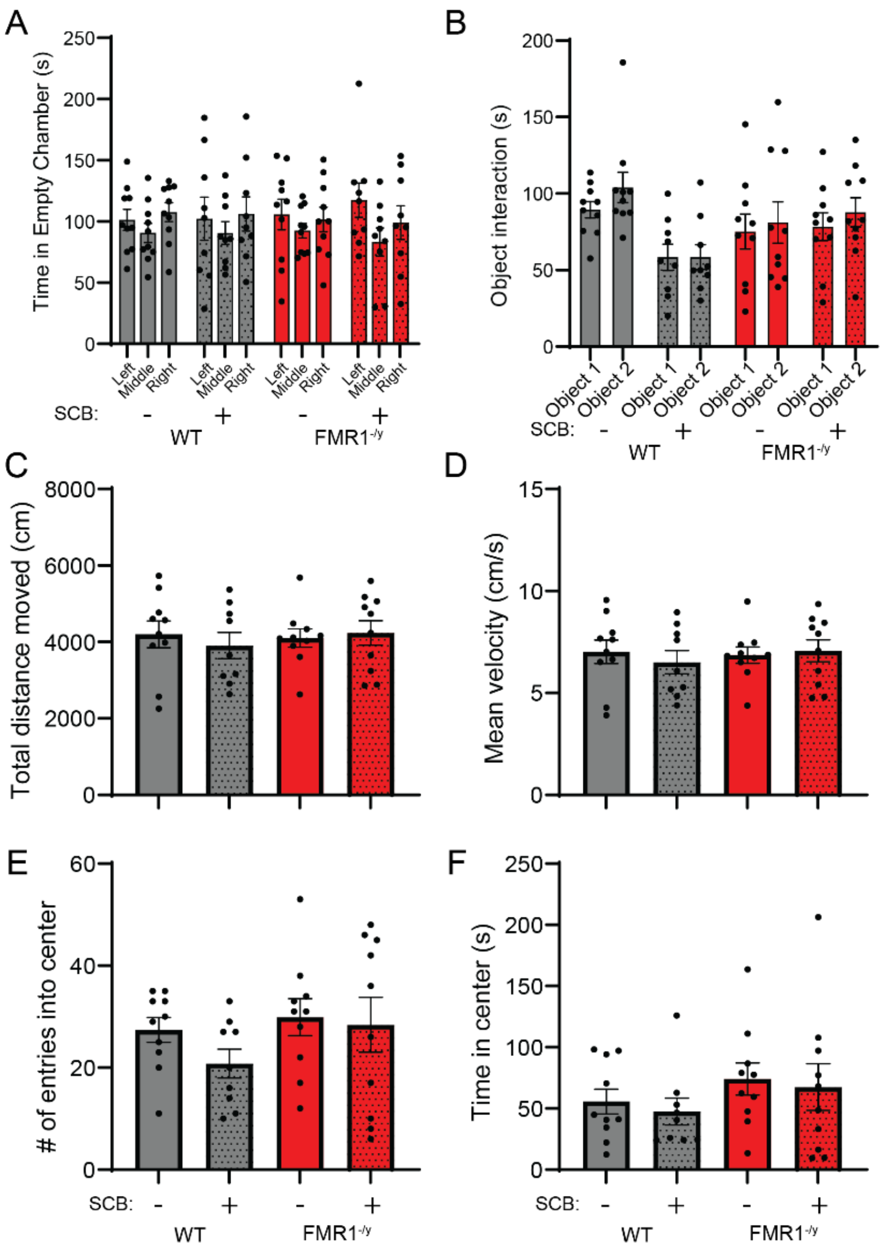


**Fig. S6. Supplemental data for behavioral tests.** A) Time spent in each of the three chambers of the social test during the acclimation period reveals no innate preference. The 'social' and 'non-social' sides were counterbalanced between left and right. B) Time spent interacting with object 1 vs object 2 on day 1 of the object recognition test reveals no innate preference for object location. The location of the 'novel' object on day 2 was counterbalanced. C-F) Performance in the open field test. No group differences were observed when total distance moved (C), overall mean velocity (D), number of entries into the center (E), and time spent in the center (F) were measured. For all tests, N=9-10 mice derived from 5 litters per condition.

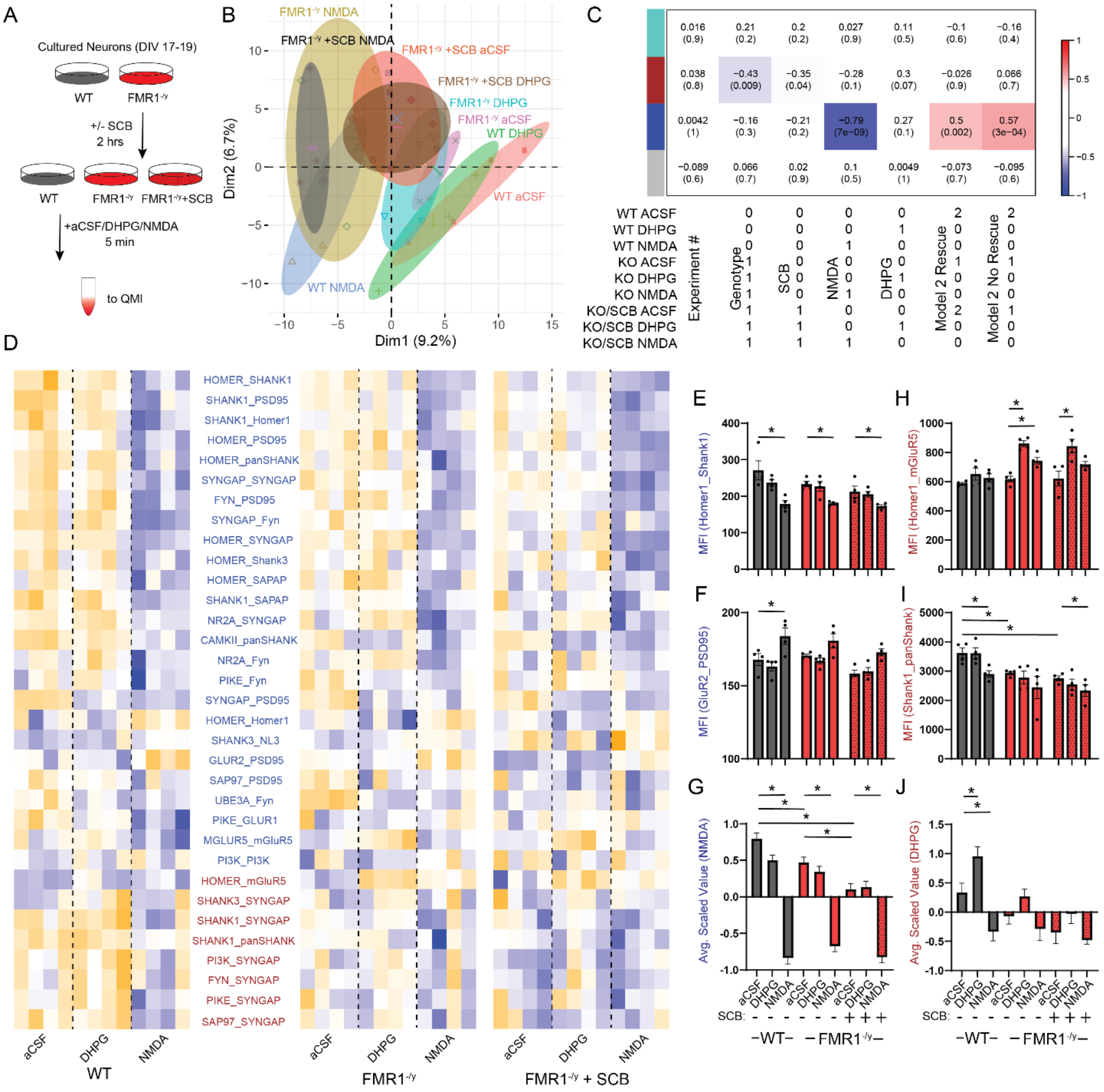


**Figure S7: SCB does not normalize basal hyperactivation of gsPIN interactions in cultured neurons.** A) Experimental design. B) Principal component graph. C) Module-trait correlation table shows the correlation coefficient (top number) and p-value (bottom number) of the correlation between each module eigenvector and the binary-coded hypothesis listed below the table. D) Interactions that were both individually statistically significant by a Bonferroni-corrected adaptive non-parametric test corrected for multiple comparisons (ANC, see methods) comparing treatment groups, and were a member of a significant module, are represented by a row-scaled heatmap (blue- low abundance, orange- high abundance). Interactions are colored based on their assigned module. E,F) Example of interactions in the NMDA-responsive blue module, Homer1_Shank1 (E) and GluR2_PSD95 (F). MFI, median fluorescent intensity. Asterisk represents statistically significant difference measured by ANC. G) Averaged scaled value of all interactions in the NMDA-responsive blue module. Asterisks represent statistically significant difference by one-way ANOVA followed by Sidak's multiple comparison test,

$p<0.05$. H,I) Examples of interactions in the genotype-correlated brown module, Homer1_mGLuR5 (H) and Shank1_panShank (I). Asterisks represents statistically significant difference by ANC, $p<0.05$. J) Averaged scaled value of all interactions in the brown module. Asterisks represent statistically significant difference by one-way ANOVA followed by Sidak's multiple comparison test, $p<0.05$. N = 4 cultures derived from 4 individual litters per condition.

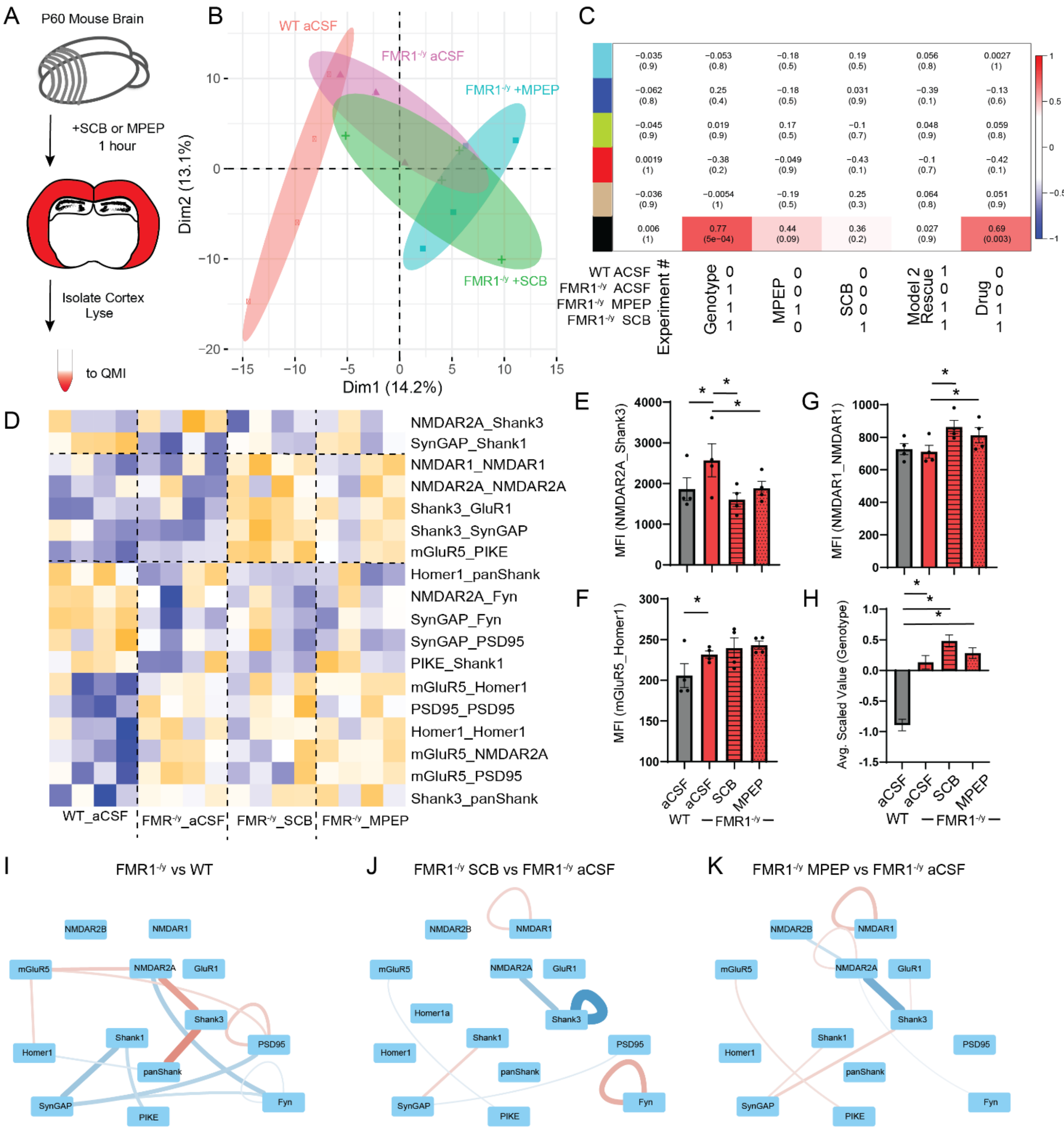

**Figure S8: SCB does not normalize protein interaction networks in acute slices from adult animals.** A) Experimental design. B) Principal component graph. C) Module-trait correlation table shows the correlation coefficient (top number) and p-value (bottom number) of the correlation between each module eigenvector and the binary-coded hypothesis listed below the table. D) Interactions that were both individually statistically significant by a Bonferroni-corrected adaptive non-parametric test corrected for multiple comparisons (ANC, see methods) comparing treatment groups, and were a member of a significant module, are represented by a row-scaled heatmap (blue- low abundance, orange- high abundance). Interactions are colored based on their assigned module. E) Two interactions were ‘normalized’ by SCB and MPEP treatment, including NMDAR2A_Shank3. MFI, median fluorescent intensity. F) Example of an interaction that was abnormal in $FMR1^{-/y}$ animals but was not altered by drug treatment, mGluR5_Homer1. G) Example of an interaction that changed with SCB/MPEP treatment, NMDAR1_NMDAR1. Asterisks represents statistically significant difference by ANC, $p<0.05$. N = 4 WT and 4 $FMR1^{-/y}$ animals; slices from each $FMR1^{-/y}$ animal were treated with aCSF, SCB and MPEP to directly compare the effects of treatment. H) Averaged scaled value of all interactions in the genotype-correlated black module. Asterisks represent statistically significant difference by one-way ANOVA followed by Sidak’s multiple comparison test, $p<0.05$. I-K) Node-edge diagrams show protein interactions were significantly different between untreated $FMR1^{-/y}$ and WT slices (I), $FMR1^{-/y}$ SCB-treated and $FMR1^{-/y}$ untreated slices (J), or $FMR1^{-/y}$ MPEP-treated and $FMR1^{-/y}$ untreated slices (K). Nodes represent proteins, edges represent protein-protein interactions that were significantly different in the indicated comparison; red = increased, blue = decreased. The relative magnitude of the change is reflected in edge thickness. Statistical significance was assessed by a Bonferroni-corrected adaptive non-parametric test corrected for multiple comparisons (ANC). N = 4 animals per condition.

**Table S1: IP probe antibody panels used for QMI**

Scaffold and structural proteins

| Target | QMI Component | Clone | Supplier | Cat# | RRID |
|---|---|---|---|---|---|
| Homer1 | IP<br>Probe | AT1F3<br>D3 | LifeSpan Bioscience<br>Santa Cruz | LS-C103482<br>sc-17842 | AB_2264378<br>AB_627742 |
| Homer1a | IP<br>Probe | NA<br>Polym13 | NA<br>Santa Cruz | NA<br>sc-8922 | NA<br>AB_648368 |
| NL3 | IP<br>Probe | 566209<br>G2 | ThermoFisher<br>Santa Cruz | MA5-24253<br>sc-271880 | AB_2576897<br>AB_10709173 |
| panShank | IP<br>Probe | NA<br>N23B/49 | NA<br>Neuromab | NA<br>75-089 | NA<br>AB_10672418 |
| PSD-95 | IP<br>Probe | K28/74<br>K28/43 | Biolegend<br>Biolegend | 810301<br>810401 | AB_2564749<br>AB_2564750 |
| SAPAP | IP<br>Probe | NA<br>N127/31 | NA<br>Biolegend | NA<br>832601 | NA<br>AB_2564958 |
| SAP97 | IP<br>Probe | RPI197.4<br>polyh60 | Enzo Life Sciences<br>Santa Cruz | ADI-VAM-PS00<br>sc-25661 | AB_1083921<br>AB_2092029 |
| Shank1 | IP<br>Probe | N22/21<br>N22/21 | Neuromab<br>Neuromab | 75-064<br>75-064 | AB_2270283<br>AB_2270283 |
| Shank3 | IP<br>Probe | N367/62<br>N69/46 | Neuromab<br>Neuromab | 75-344<br>75-109 | AB_2315921<br>AB_2187730 |

Receptor Proteins

| Target | QMI Component | Clone | Supplier | Cat# | RRID |
|---|---|---|---|---|---|
| GluR1 | IP<br>Probe | N355/1<br>poly1594 | Biolegend<br>Millipore | 819801<br>AB1504 | AB_2564834<br>AB_2113602 |
| GluR2 | IP<br>Probe | L21/32<br>polyc20 | Biolegend<br>Santa Cruz | 810501<br>sc-7610 | AB_2564751<br>AB_2247873 |
| mGluR5 | IP<br>Probe | 5675<br>N75/3 | Millipore<br>Neuromab | AB5675<br>75-115 | AB_2295173<br>AB_10672260 |
| NMDAR1 | IP<br>Probe | 54.1<br>polyc20 | ThermoFisher<br>Santa Cruz | 32-0500<br>sc-1467 | AB_2533060<br>AB_670215 |
| NMDAR2A | IP<br>Probe | N327/95<br>N3278/38 | Neuromab<br>Biolegend | 75-288<br>832401 | AB_2315842<br>AB_2564956 |
| NMDAR2B | IP<br>Probe | N59/20<br>N59/36 | Biolegend<br>Biolegend | 832501<br>818701 | AB_2564957<br>AB_2564823 |

Signaling proteins

| Target | QMI Component | Clone | Supplier | Cat# | RRID |
|---|---|---|---|---|---|
| CaMKII | IP<br>Probe | 6G9<br>polyC6970 | ThermoFisher<br>Sigma | MA1-048<br>C6974 | AB_325403<br>AB_258984 |
| Fyn | IP<br>Probe | Fyn15<br>Fyn59 | Santa Cruz<br>Biolegend | sc-434<br>626502 | AB_627642<br>RRID:AB_2278824 |
| PI3K p85 | IP<br>Probe | U5<br>AB6 | ThermoFisher<br>Millipore | MA1-74183<br>05-212 | AB_2163452<br>AB_309658 |
| PIKE | IP<br>Probe | 263A<br>DN8 | Bethyl Laboratories<br>Rockland | A304-263A<br>200-401-DN8 | AB_2620459<br>AB_2612161 |
| SynGap | IP<br>Probe | D20C7<br>PolyR19 | Cell Signaling<br>Santa Cruz | 5539<br>sc-8572 | AB_10694401<br>AB_2200750 |
| Ube3a | IP<br>Probe | H182<br>E4 | Santa Cruz<br>Santa Cruz | sc-25509<br>sc-166689 | AB_639982<br>AB_2211807 |
